# A hierarchy of linguistic predictions during natural language comprehension

**DOI:** 10.1101/2020.12.03.410399

**Authors:** Micha Heilbron, Kristijan Armeni, Jan-Mathijs Schoffelen, Peter Hagoort, Floris P. de Lange

## Abstract

Understanding spoken language requires transforming ambiguous acoustic streams into a hierarchy of representations, from phonemes to meaning. It has been suggested that the brain uses prediction to guide the interpretation of incoming input. However, the role of prediction in language processing remains disputed, with disagreement about both the ubiquity and representational nature of predictions. Here, we address both issues by analysing brain recordings of participants listening to audiobooks, and using a deep neural network (GPT-2) to precisely quantify contextual predictions. First, we establish that brain responses to words are modulated by ubiquitous, probabilistic predictions. Next, we disentangle model-based predictions into distinct dimensions, revealing dissociable signatures of syntactic, phonemic and semantic predictions. Finally, we show that high-level (word) predictions inform low-level (phoneme) predictions, supporting hierarchical predictive processing. Together, these results underscore the ubiquity of prediction in language processing, showing that the brain spontaneously predicts upcoming language at multiple levels of abstraction.

## Introduction

Understanding spoken language requires transforming ambiguous stimulus streams into a hierarchy of increasingly abstract representations, ranging from speech sounds to meaning. It is often argued that during this process, the brain relies on prediction to guide the interpretation of incoming information [1, 2]. Such a ‘predictive processing’ strategy has not only proven effective for artificial systems processing language [3, 4], but has also been found to occur in neural systems in related domains such as perception and motor control and might constitute a canonical neural computation [5, 6].

There is a considerable amount of evidence that appears in line with predictive language processing. For instance, behavioural and brain responses are highly sensitive to violations of linguistic regularities [7, 8] and to deviations from linguistic expectations more broadly [9–13]. While such effects are well-documented, two important questions about the role of prediction in language processing remain unresolved [14].

The first question concerns the *ubiquity* of prediction. While some models cast prediction as a routine, integral part of language processing [1, 15, 16], others view it as relatively rare, pointing out that apparent widespread prediction effects might instead reflect other processes like semantic integration difficulty [17, 18]; or that such prediction effects might be exaggerated by the use of artificial, prediction-encouraging experiments focussing on highly predictable ‘target’ words [17, 19]. The second question concerns the representational nature of predictions: Does linguistic prediction occur primarily at the level of syntax [15, 20–22] or rather at the lexical [16, 23], semantic [24, 25] or the phonological level [13, 26–29]? ERP studies have described brain responses to violations of, and deviations from, both high and low-level expectations, suggesting prediction might occur at all levels simultaneously [1, 19], although see [30]. However, it has been disputed whether these findings would generalise to natural language, where violations are rare or absent and with few highly predictable words. In these cases, prediction may be less relevant or might perhaps be limited to the most abstract levels [17, 19, 30].

Here, we address both issues, probing the ubiquity and nature of linguistic prediction during natural language understanding. Specifically, we analysed brain recordings from two independent experiments of participants listening to audiobooks, and use a state-of-the-art deep neural network (GPT-2) to quantify linguistic predictions in a fine-grained, contextual fashion. First, we obtain evidence for predictive processing, confirming that brain responses to words are modulated by *probabilistic* predictions. Critically, the effects of prediction were found over and above those of non-predictive factors such as integration difficulty, and were not confined to a subset of predictable words, but were widespread – supporting the notion of *ubiquitous* prediction. Next, we investigated at which level prediction occurs. To this end, we disentangled the model-based predictions into distinct dimensions, revealing dissociable neural signatures of syntactic, phonemic and semantic predictions. Finally, we found that higher-level (word) predictions constrain lower-level (phoneme) predictions, supporting hierarchical prediction. Together, these results underscore the ubiquity of prediction in language processing, and demonstrate that prediction is not confined to a a single level of abstraction but occurs throughout the language network, forming a hierarchy of predictions across all levels of analysis, from phonemes to meaning.

## Results

We consider data from two independent experiments, in which brain activity was recorded while participants listened to natural speech from audiobooks. The first experiment is part of a publicly available dataset [31], and contains 1 hour of electroencephalographic (EEG) recordings in 19 participants. The second experiment collected 9 hours of magneto-encephalographic (MEG) data in three individuals, using individualised head casts that allowed us to localise the neural activity with high precision. While both experiments had a similar setup (see Figure 1), they yield complementary insights, both at the group level and in three individuals.

**Figure 1:**
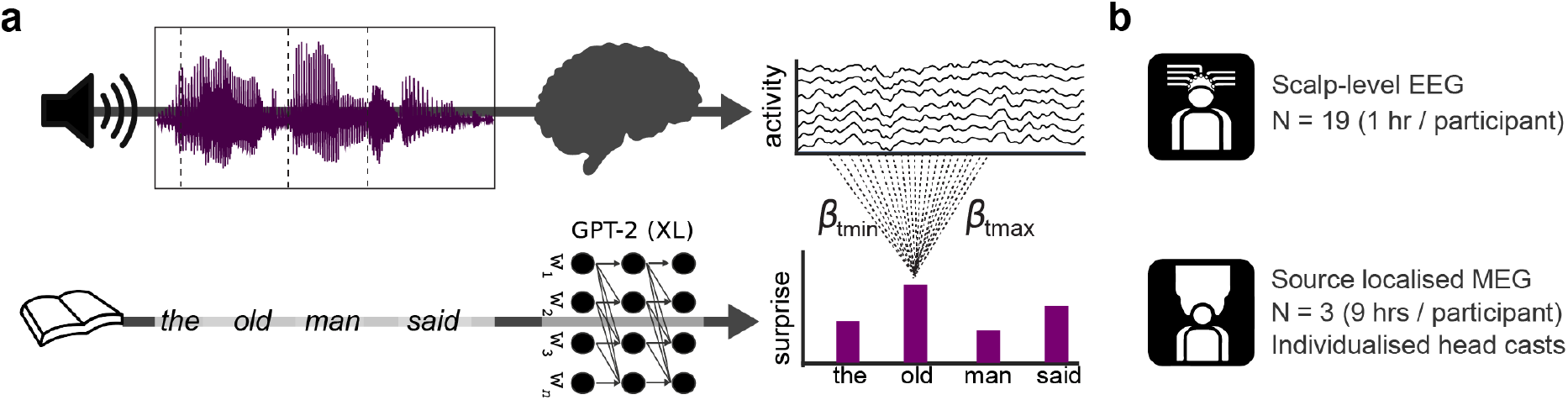
Schematic of experimental and analytical framework. **a)** Top row: in both experiments participants listened to continuous recordings from audiobooks while brain activity was recorded. Bottom row: the texts participants listened to were analysed by a deep neural network (GPT-2) to quantify the contextual probability of each word. A regression-based technique was used to estimate the effects of (different levels of) linguistic unexpectedness on the evoked responses within the continuous recordings. **b)** Datasets analysed: one group-level EEG dataset, and one individual subject source-localised MEG dataset.

### Neural responses to speech are modulated by probabilistic linguistic predictions

We first tested for evidence for linguistic prediction in general. We reasoned that if the brain is constantly predicting upcoming language, neural responses to words should be sensitive to violations of contextual predictions, yielding ‘prediction error’ signals which are considered a hallmark of predictive processing [5]. To this end, we used a regression-based deconvolution approach to estimate the effects of prediction error on evoked responses within the continuous recordings. We focus on this event-related, low-frequency evoked response because it connects most directly to earlier influential neural signatures of prediction in language [7, 30, 32, 33].

To quantify linguistic predictions, we analysed the books participants listened to with a state-of-the-art neural language model: GPT-2 [34]. GPT-2 is a large transformer-based model that predicts the next word given the previous words, and is currently among the best publicly-available models of its kind. Note that we do not use GPT-2 as a model of human language processing, but purely as a tool to quantify how expected each word is in context.

To test whether neural responses to words are modulated by contextual predictions, we compared three regression models (see S5). The baseline model formalises the hypothesis that natural, passive language comprehension does not invoke prediction. This model did not include regressors related to contextual predictions, but did include several potentially confounding variables (such as word frequency, semantic integration, and acoustics). The *constrained guessing* model formalised the hypothesis that language processing *sometimes* (in constraining contexts) invokes prediction, and that such predictions are an all-or-none phenomenon – together representing how the notion of prediction was classically used in the psycholinguistic literature [33]. This model included all non-predictive variables from the baseline model, plus, in constraining contexts, a linear estimate of word improbability (since all-or-none predictions result in a linear relationship between word probability and brain responses; see methods for details). Finally, the *probabilistic prediction* model included all confounding regressors from the baseline model, plus for every word a logarithmic estimate of word improbability (i.e. *surprise*). This formalises the hypothesis that the brain constantly generates *probabilistic predictions*, as proposed by predictive processing accounts of language [1, 32] and of neural processing more broadly [5, 6].

When we compared the ability of these models to predict brain activity using cross-validation, we found that the probabilistic prediction model performed better than both other models (see Figure 2a). The effect was highly consistent, found in virtually all EEG participants (probabilistic vs constrained guessing, *t*_18_ = 5.34, *p* = 4.46 × 10^−5^; probabilistic vs baseline, *t*_18_ = 6.43, *p* = 4.70 × 10^−6^) and within each MEG participant (probabilistic vs constrained guessing, all *p*^′^*s <* 1.54 × 10^−6^; probabilistic vs baseline, all *p*^′^*s <* 5.17 × 10^−12^).

**Figure 2:**
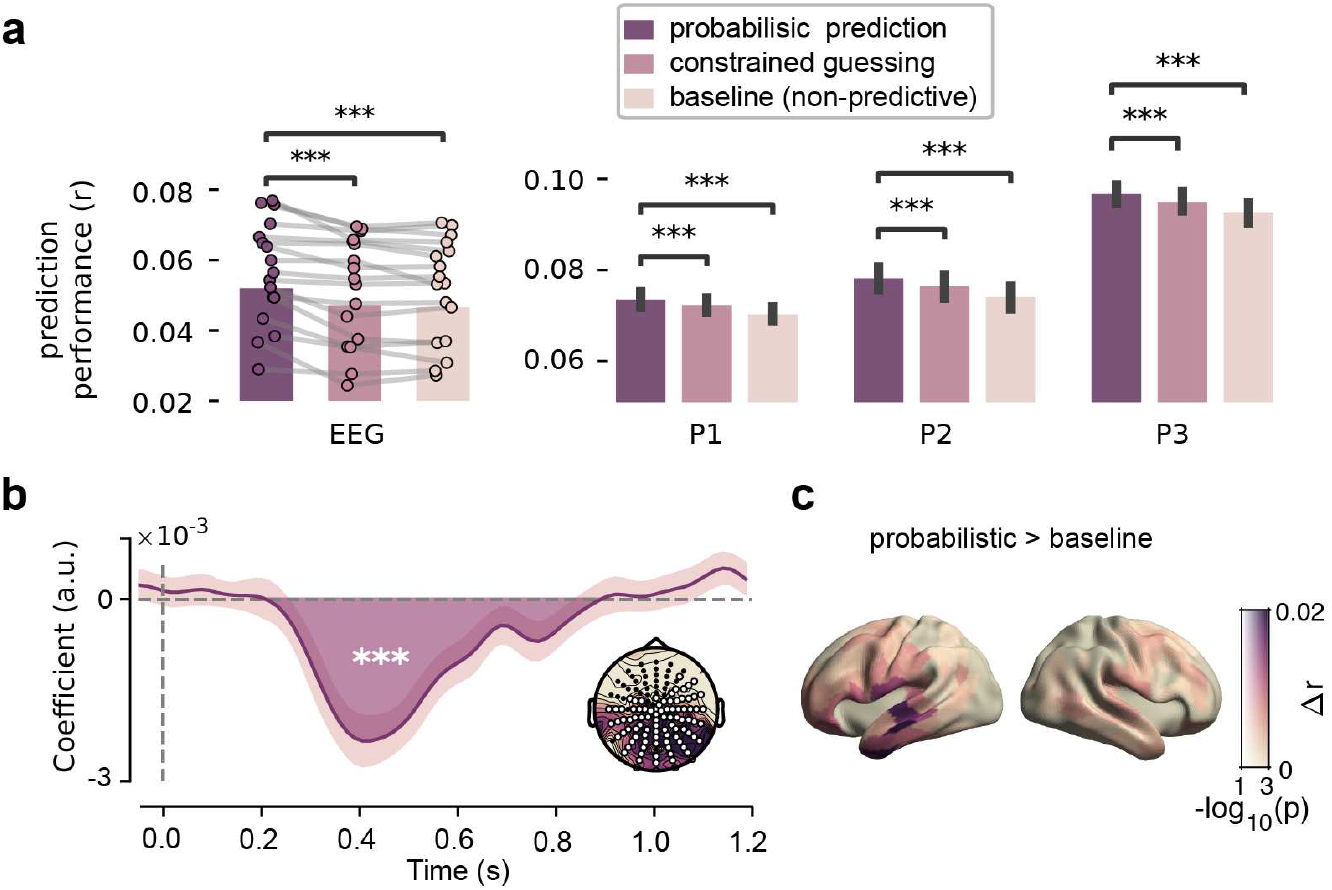
Neural responses are modulated by probabilistic predictions. **a)** Model comparison. Cross-validated correlation coefficients for EEG (left) and each MEG participant (right). EEG: dots with connecting lines represent individual participants (averaged over all channels). MEG: bars represent median across runs, bars represent bootstrapped absolute deviance (averaged over language network sources). **b)** EEG: coefficients describing the significant effect of lexical surprise (see Figure S3 for the full topography over time). Highlighted area indicates extent of the cluster, shaded error bar indicates bootstrapped SE. Inset shows distribution of absolute t-values and of channels in the cluster. **c)** Difference in prediction performance across cortex (transparency indicates FWE-corrected p-values). Significance levels correspond to P<0.001 (***) in a two-tailed one-sample Student’s *t* or Wilcoxon sign rank test.

As the *constrained guessing* model differed from the probabilistic model in two ways – by assuming that predictions are (i) categorical and (ii) limited to constraining contexts – we also considered a control model. Like the constrained guessing model, this extended guessing model included a linear estimate of word probability, but for every word rather than only for constraining contexts. Although this model did not outperform the probabilistic prediction model, it did substantially outperform the constrained model (Fig S5). This demonstrates that the effects of prediction are not limited to constraining contexts, but apply much more broadly – in line with the idea that predictions are ubiquitous and automatic.

Having established that word unexpectedness modulates neural responses, we characterised this effect in space and time. In the MEG dataset, we asked for which neural sources lexical surprise was most important in explaining neural data, by comparing the prediction performance of the baseline model to the predictive model in a spatially resolved manner. This revealed that overall word unexpectedness modulated neural responses throughout the language network (see Figure 2c). To investigate the temporal dynamics of this effect, we inspected the regression coefficients, which describe how fluctuations in lexical surprise modulate the neural response at different time lags – together forming a modulation function also known as the *regression evoked response* [35] or Temporal Response Function (TRF) [27, 36]. When we compared these across participants in the EEG experiment, cluster-based permutation tests revealed a significant effect (*p* = 2 × 10^−4^) based on a posterio-central cluster with a negative polarity between 0.2 and 0.9 seconds (see Figure 2b and S8). This indicates that surprising words lead to a stronger negative deflection of evoked responses, an effect peaking at 400 ms post word onset and strongly reminiscent of the classic N400 [7, 24, 30]. Coefficients for MEG subjects revealed a similar, slow effect at approximately the same latencies (see Fig S4).

Together, these results constitute clear evidence for predictive processing by confirming that brain responses to words are modulated by predictions. These modulations are not confined to constraining contexts, occur throughout the language network, evoke an effect reminiscent of the N400, and are best explained by a probabilistic account of prediction. This suggests the brain predicts constantly and probabilistically – even when passively listening to natural language.

### Linguistic predictions are feature-specific

The results so far revealed modulations of neural responses by *overall* word unexpectedness. What type of linguistic prediction might be driving these effects? Earlier research suggests a range of possibilities, with some proposing that the effect of overall word surprise primarily reflects syntax [15, 20], while others propose that prediction unfolds at the seman-tic [24, 25], or the phonemic level [13, 26, 27] – or at all levels simultaneously [1].

To evaluate these possibilities, we factorised the aggregate, word-level linguistic predictions from the artificial neural network into distinct linguistic dimensions (Fig 3). This allows us to derive model-based estimates of three feature-specific predictions: the syntactic prediction (defined as the conditional probability distribution over parts-of-speech, given context), semantic prediction (defined as the predicted semantic embedding) and phonemic prediction (i.e. the conditional probability of the next phoneme, given the phonemes within the word so far and the prior context). By comparing these predictions to the presented words, we derived *feature-specific prediction errors* which quantified not just the extent to which a word is surprising overall, but also in what way: semantically, syntactically or phonemically (see Methods for definitions).

**Figure 3:**
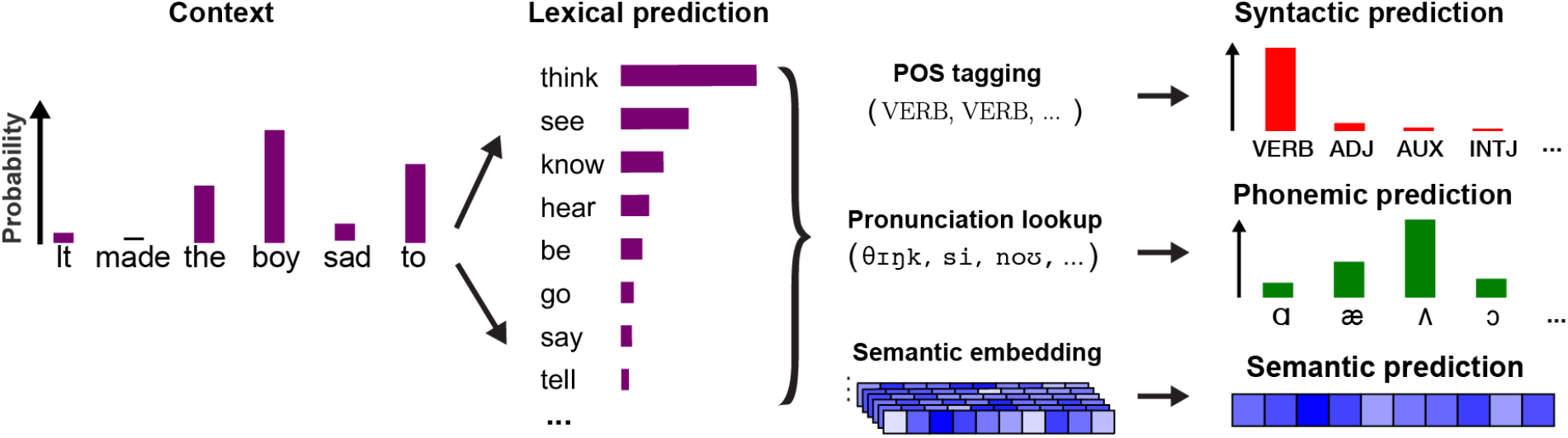
Partitioning model-derived predictions into distinct linguistic dimensions. To disentangle syntactic, semantic and phonemic predictions, the lexical predictions from GPT-2 were analysed. For the syntactic prediction, part-of-speech was tagging performed over all potential sentences (e.g. “It made the boy sad to *think*”). To compute the phonemic prediction, each predicted word was decomposed into its constituent phonemes, and the predicted probabilities were used as a contextual prior in a phoneme model (see Figure 6). For the semantic prediction, a weighted average was computed over the GLoVE embeddings of all predicted words.

We reasoned that if the brain is generating predictions at a given level (e.g. syntax), then the neural responses should be sensitive to prediction errors specific to this level. Moreover, because these different features are processed by partly different brain areas over different timescales, the prediction errors should be at least partially dissociable. To test this, we formulated a new regression model (Figure S6). This included all variables from the lexical prediction model as nuisance regressors, and added three regressors of interest: syntactic surprise (defined for each word), semantic prediction error (defined for each content word), and phonemic surprise (defined for each word-non-initial phoneme).

Because these regressors were to some degree correlated, we first asked whether, and in which brain area, each of the feature-specific prediction errors explained any unique variance, not explained by the other regressors. In this analysis, we turn to the MEG data because of its spatial specificity. As a control, we first performed the analysis for a predictor with a known source: the acoustics. This revealed a clear peak around auditory cortex (Fig S7) especially in the right hemisphere. This aligns with prior work [37] and confirms that this approach can localise which areas are especially sensitive to a given regressor. We then tested the three prediction errors, finding that each type of prediction error explained significant unique variance in each individual (Figure 4), except in participant 1 where phonemic surprise did not survive multiple comparisons correction (but see Figure 6c and Discussion). This shows that the brain responds differently to different types of prediction errors, implying that linguistic predictions are feature-specific and occur both at high and low levels of processing simultaneously.

**Figure 4:**
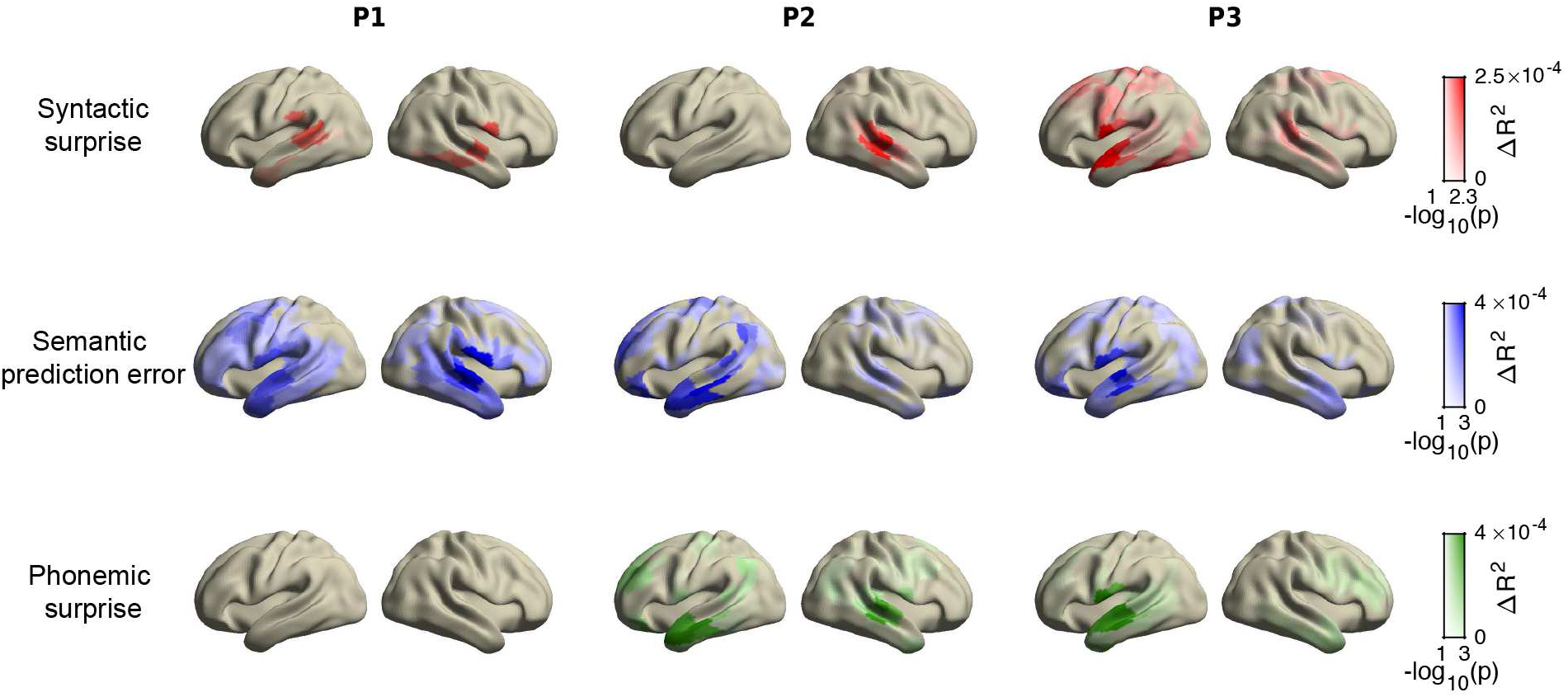
Dissociable patterns of explained variance by syntactic, semantic and phonemic predictions. Unique variance explained by syntactic, semantic and phonemic unexpectedness (quantified via surprise or prediction error) across cortical sources in each MEG participant. In all plots, colour indicates amount of additional variance explained; opacity indicates FWE-corrected statistical significance. Note that *p <* 0.05 is equivalent to — log_10_(*p*) *>* 1.3.

Although we observed considerable variation in lateralisation and exact spatial locations between individuals, the overall pattern of sources aligned well with prior research on the neural circuits for each level. For instance, only for semantic prediction errors we observed a widely distributed set of neural sources – consistent with the fact that the semantic (but not the syntactic or phonological) system is widely distributed [38, 39]. Moreover, the temporal areas showing the strongest effect of syntactic surprise are indeed key areas for syntactic processing [40] and for the posterior temporal areas predictive syntax in particular [21, 41–43] – though a clear syntactic effect in the inferior frontal gyrus (IFG) was interestingly absent. When we compared the sources of phonemic surprise to those obtained for lexical surprise, we observed a striking overlap in all individuals (see Fig. S7, S4 and S13), suggesting that the phonemic predictions as formalised here mostly relate to predictive (incremental) word recognition at the phoneme level rather than describing phonological or phonotactic predictions *per se*.

### Dissociable signatures of syntactic, semantic and phonemic predictions

Having established that syntactic, phonemic and semantic prediction errors independently modulated neural responses in different brain areas, we further investigated the nature of these effects. This was done by inspecting the coefficients (or modulation functions), which describe how fluctuations in a given regressor modulate the response over time. We first turn to the EEG data because there the sample size allows for population-level statistical inference on the coefficients. We fitted the same integrated model (Figure S6) and performed cluster-based permutation tests on the modulation functions. This revealed significant effects for each type of prediction error (Figure 5).

**Figure 5:**
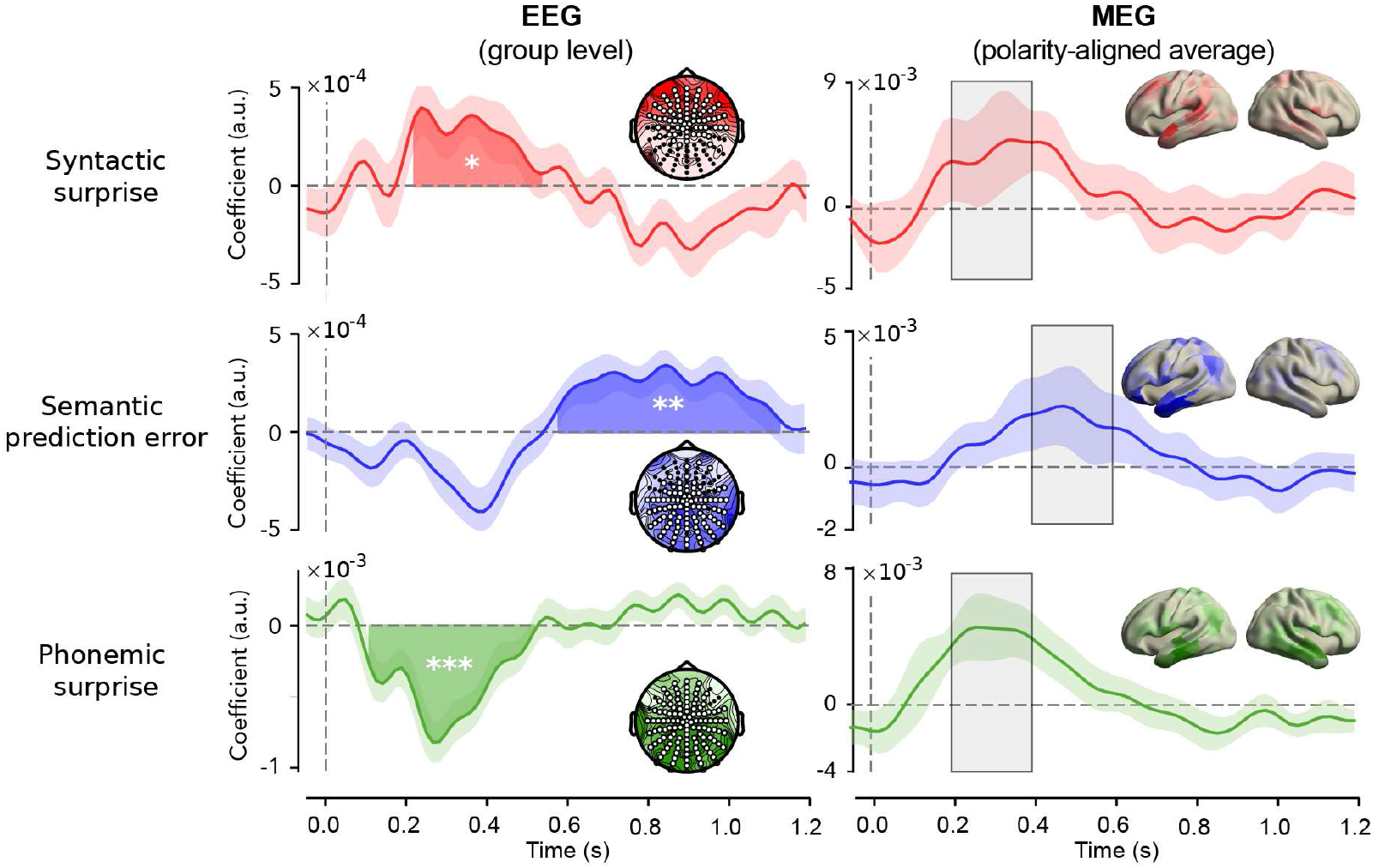
Spatiotemporal signatures of syntactic, semantic and phonemic prediction errors. Coefficients describing the effects of each prediction-error. EEG (left column): modulation functions averaged across the channels participating for at least one sample in the three main significant clusters (one per predictor). Highlighted area indicates temporal extent of the cluster. Shaded area around waveform indicates bootstrapped standard errors. Stars indicate cluster-level significance; *p <* 0.05 (*), *p <* 0.05 (**), *p <* 0.001 (***). Insets represent selected channels and distribution of absolute t-values. Note that these plots only visualise the *effects*; for the full topographies of the coefficients and respective statistics, see Figure S8. MEG (right column): polarity aligned responses averaged across the sources with significant explained variance (Figure 4) across participants. Shaded area represents absolute deviation. Insets represent topography of absolute value of coefficients averaged across the highlighted period. Note that due to polarity alignment, sign information is to be ignored for the MEG plots. For average coefficients for each source, see Figure S10; for coefficients of each individual, see Figs S11 - S14.

First, syntactic surprise evoked an early, positive deflection (*p* = 0.027) based on a frontal cluster between 200 and 500 ms. This early frontal positivity converges with two recent studies that investigated specifically syntactic prediction using models trained explicitly on syntax [22, 44]. We also observed a late negative deflection for syntactic surprise (*p* = 0.025; Figure S9), but this was neither in line with earlier findings nor replicated in the MEG data. The semantic prediction error also evoked a positive effect (*p* = 9.1 × 10^−3^) but this was based on a much later, spatially distributed cluster between 600 and 1100 ms. Although such a late positivity has been prominently associated with *syntactic* violations [8], there is also a considerable body of work reporting such late positivities for purely semantic anomalies [45] which is more in line with the semantic prediction error as quantified here (see Discussion). Notably, we did not find a significant N400-like effect for semantic prediction error – possibly because this negative deflection was already explained by the overall lexical surprise, which was included as a nuisance regressor (Figure S10). Finally, the phonemic surprise evoked a negative effect (*p* = 3 × 10^−4^) based on an early, distributed cluster between 100 and 500 ms. This effect was similar to the word-level surprise effect (Figure 2C and S10) but occurred earlier. This timecourse corresponds to recent studies using similar regression-based techniques to study (predictive) phoneme processing in natural listening [13, 28, 46].

When we performed the same analysis on the MEG data, we observed striking differences in the exact shape and timing of the modulation functions between individuals (see Figure S11-S14). While this might partly reflect variance in the coefficients due to inherent correlations between the variables, it clearly also reflects true individual differences, demonstrated by one of the strongest and least correlated regressors (the acoustics) also showing considerable variability (see Figure S14). Overall however, we could recover a temporal pattern of effects similar to the EEG results: phonemic and syntactic surprise modulating early responses, and semantic prediction error modulating later responses – although not as late in the EEG data. This temporal order holds on average (Figures 5, S10) and is especially clear within individuals (Figure S11 - S13).

Overall, our results (Figure 4,5) demonstrate that syntactic, phonemic and semantic prediction errors evoke brain responses that are both temporally and spatially dissociable. Specifically, while phonemic and syntactic predictions modulate relatively early neural responses (100-400 ms) in a set of focal temporal (and frontal) areas that are key for syntactic and phonetic/phonemic processing, semantic predictions modulate later responses (>400 ms) across a widely distributed set of areas across the distributed semantic system. These results reveal that linguistic prediction is not implemented by a single system but occurs throughout the speech and language network, forming a hierarchy of linguistic predictions across all levels of analysis.

### Phoneme predictions reveal hierarchical inference

Having established that the brain generates linguistic predictions across multiple levels of analysis, we finally asked whether predictions at different levels might interact. One option is that they are encapsulated: Predictions in separate systems might use different information, for instance unfolding over different timescales, rendering them independent. Alternatively, predictions at different levels might inform and constrain each other, effectively converging into a single multilevel prediction – as suggested by theories of hierarchical cortical prediction [5, 6,47].

One way to adjudicate between these hypotheses is by evaluating different schemes of deriving phoneme predictions. One possibility is that such predictions are only based on information unfolding over short timescales. In this scheme, the predicted probability of the next phoneme is derived from the *cohort* of words that are compatible with the phonemes presented so far, with each candidate word weighted by its overall frequency of occurrence (see Figure 6A). As such, this scheme proposes a *single-level model*: phoneme predictions are based only on information at the level of within-word phoneme sequences unfolding over short timescales, plus a fixed frequency-based prior (capturing statistical knowledge of word frequencies within a language).

**Figure 6:**
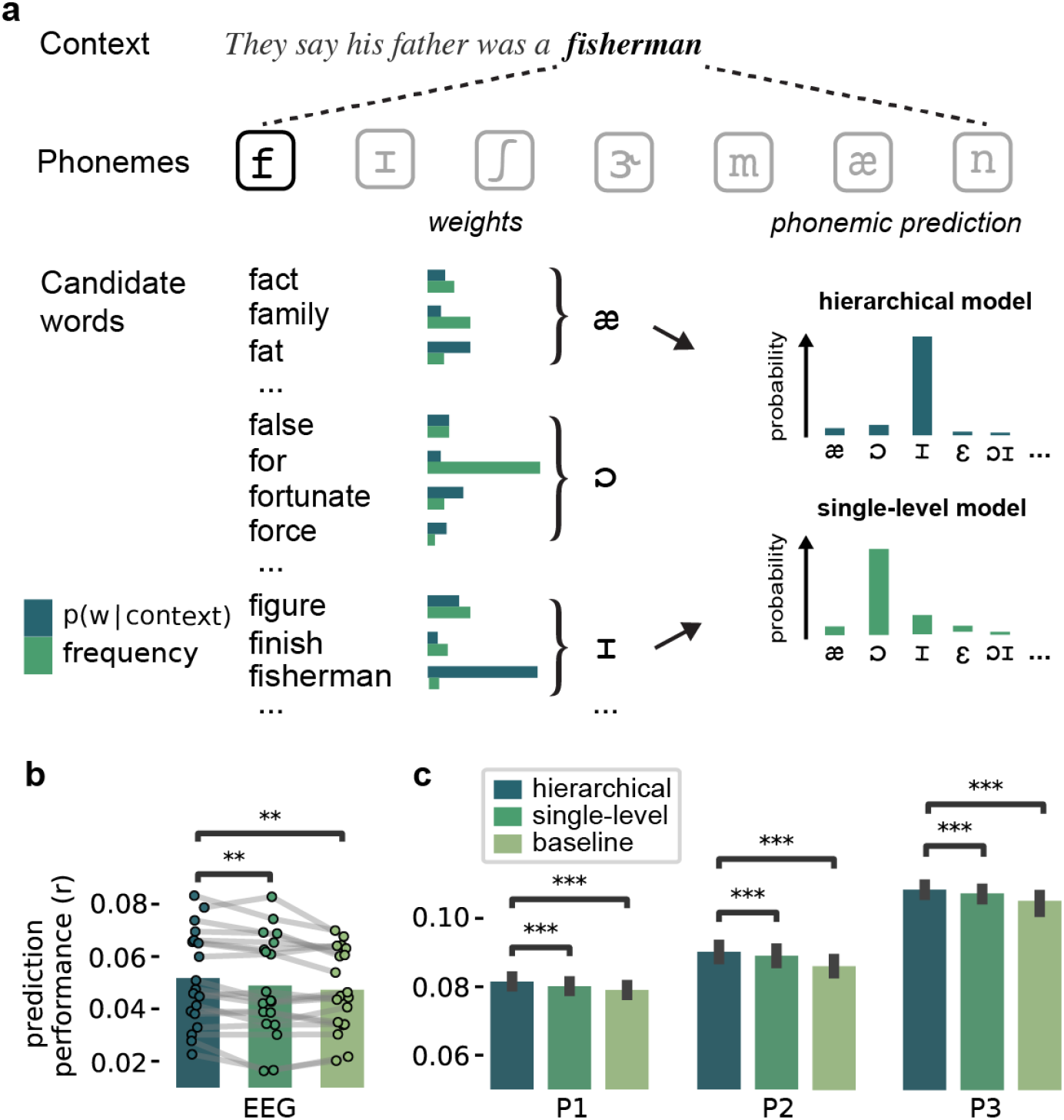
Evidence for hierarchical inference during phoneme prediction. **a)** Two models of phoneme prediction during incremental word recognition. Phonemic predictions were computed by grouping candidate words by their identifying next phoneme, and weighting each candidate word by its prior probability. This weight (or prior) could be either based on a word’s overall probability of occurrence (i.e. frequency) or on its conditional probability in that context (from GPT-2). Critically, in the frequency-based model, phoneme predictions are based on a single level: short sequences of within words phonemes (hundreds of ms long) plus a fixed prior. By contrast, in the contextual model, predictions are based not just on short sequences of phonemes, but also on a contextual prior which is itself based on long sequences of prior words (up to minutes long), rendering the model hierarchical (see Methods). **b-c)** Model comparison results in EEG (**b**) and all MEG participants (**c**). EEG: dots with connecting lines represent individual participants (averaged over all channels). MEG: bars represent median across runs, error bars represent bootstrapped absolute deviance (averaged over language network sources). Significance levels correspond to P<0.01 (**) or P<0.001 (***) in a two-tailed paired *t* or Wilcoxon sign rank test.

Alternatively, phoneme predictions might not only be based on sequences of phonemes within a word, but also on the longer prior linguistic context. In this case, the probability of the next phoneme would still be derived from the cohort of words compatible with the phonemes presented so far, but now each candidate word is not weighted by its overall frequency but by its *contextual probability* (Figure 6A). Such a model would be hierarchical, in the sense that predictions are based both – at the first level – on short sequences of phonemes (i.e. of hundreds of milliseconds long), and on a contextual prior which itself is based – at the higher level – on long sequences of words (i.e. of tens of seconds to minutes long).

Here, the first model is more in line with the classic Cohort model of incremental (predictive) word recognition, which suggests that context is only integrated after the selection and activation of lexical candidates [48]. By contrast, the second model is more in line with contemporary theories of hierarchical predictive processing which propose that high-level cortical predictions (spanning larger spatial or temporal scales) inform and shape low-level predictions (spanning finer spatial or temporal scales) [47, 49]. Interestingly, recent studies of phoneme predictions during natural listening have used both the frequency-based single level model [27, 29] and a context-based (hierarchical) model [13]. However, the models have not been explicitly compared to test which model can best account for prediction-related fluctuations in neural responses to phonemes.

To compare these possibilities, we constructed 3 phoneme-level regression models (see Figure S15), which all only included regressors at the level of phonemes. First, the baseline model only included non-predictive control variables: phoneme onsets, acoustics, word boundaries and uniqueness points. This can be seen as the phoneme-level equivalent of the baseline model in Figure **??**. The baseline model was compared with two regression models which additionally included phoneme surprise. In one of the regression models, this was calculated using a single-level model (with a fixed, frequency-based prior), in the other regression model it was derived from a hierarchical model (with a dynamic, contextual prior derived from GPT-2). To improve our ability to discriminate between the hierarchical and single-level model, we not only included surprise but also phoneme entropy (calculated with either model) as a regressor [13].

When we compared the cross-validated predictive performance, we first found that in both datasets the predictive model performed significantly better than the non-predictive baseline (Figure 6b-c hierarchical vs baseline, EEG: *t*_18_ = 3.80, *p* = 1.31 × 10^−3^; MEG: all *p*^′^*s <* 5.69 × 10^−12^). This replicates the basic evidence for predictive processing but now at the phoneme rather than word level (Figure **??**). Critically, when we compared the two predictive models, we found that the hierarchical model performed significantly better, both in EEG (*t*_18_ = 3.03, *p* = 7.28 × 10^−3^) and MEG (all *p*^′^*s <* 9.44 × 10^−4^). This suggests that neural predictions of phonemes (based on short sequences of within-word speech sounds) are are informed by lexical predictions, effectively incorporating long sequences of prior words as contexts. This is a signature of hierarchical prediction, supporting theories of hierarchical predictive processing.

## Discussion

Across two independent data sets, we combined deep neural language modelling with regression-based deconvolution of human electrophysiological (EEG and MEG) recordings to ask if and how evoked responses to speech are modulated by linguistic expectations that arise naturally while listening to a story. Our results demonstrated that evoked responses are modulated by *probabilistic* predictions. We then introduced a novel technique that allowed us to quantify not just how much a linguistic stimulus is surprising, but also at what level – phonemically, syntactically and/or semantically. This revealed dissociable effects, in space and time, of different types of prediction errors: syntactic and phonemic prediction errors modulated early responses in a set of focal, mostly temporal areas, while semantic prediction errors modulated later responses across a widely distributed set of cortical areas. Finally, we found that phonemic prediction error signals were best modelled by a hierarchical model incorporating two levels of context: short sequences of within-word phonemes (up to hundreds of milliseconds long) and long sequences of prior words (up to minutes long). Together, these results demonstrate that during natural listening, the brain is engaged in prediction across multiple levels of linguistic representation, from speech sounds to meaning. The findings underscore the ubiquity of prediction during language processing, and fit naturally in predictive processing accounts of language [1, 2] and neural computation more broadly [5, 6, 49, 50].

A primary result of this paper is that evoked responses to words are best explained by a predictive processing model: regression models including unexpectedness performed better than strong non-predictive baseline models, demonstrating that the effects of prediction on brain responses cannot be reduced to confounding simple features like semantic incongruency. This aligns with recent ERP studies aimed specifically at distinguishing prediction from semantic integration [51, 52] and extends those findings by analysing not just specific (highly predictable) ‘target’ words, but *all* words in a natural story. Indeed, when we further compared different accounts of prediction, responses were best explained by a regression model casting linguistic predictions as ubiquitous and probabilistic. This supports the notion of continuous, graded prediction – as opposed to the classical view of prediction as the all-or-none pre-activation of specific words in highly constraining contexts [33].

Because our deconvolution analysis focussed on evoked responses, the results can be linked to the rich literature on linguistic violations using traditional ERP methods. This is powerfully illustrated by the modulation function of lexical surprise (Figure 2b) tightly following the N400 modulation effect, one of the first proposed, most robust and most debated ERP signatures of linguistic prediction [7, 24, 30]. Similarly, the early negativity we found for phonemic surprise and later positivity for semantic prediction error (Fig 5) align well with N200 and the semantic P600 or PNP effects of phonological mismatch and semantic anomaly respectively [33, 53]. Unlike most ERP studies, we observed these effects in participants listening to natural stimuli – without any anomalies or violations – not engaged in any task. This critically supports the idea that these responses reflect deviations from *predictions* inherent to the comprehension process – rather than reflecting either detection of linguistic anomalies or expectancy effects introduced by the experiment [17, 19].

While we found several striking correspondences between the modulation functions recovered from the data and classic effects from the ERP literature, there were also some differences. Specifically, for syntactic surprise, we found neither a late positive effect resembling the syntactic P600 [8] nor an early negative effect akin to the ELAN [54]. One potential explanation for this is that our formalisation (part-of-speech surprise) might not fully capture syntactic violations used in ERP studies. Indeed, a recent paper on syntactic prediction using a similar model-based approach found a P600-like effect not for syntactic surprise but for the number of syntactic reinterpretation attempts a word induced [22]. Conversely, the early positive effect of syntactic surprise we found – which replicated other model-based findings, despite using a different formalisation of syntactic surprise [22, 44] – does not have a clear counterpart in the traditional ERP literature. Better understanding such systematic differences between the traditional experimental and model-based approach provides an interesting challenge for future work.

Beyond the ERP literature, there has also been earlier model-based work on prediction. However, these studies have mostly quantified feature-unspecific lexical unexpectedness [10, 12, 32, 55, 56] or modelled feature-specific predictions at a single level such as syntax [11, 22, 44, 57], phonemes [13, 27, 28] or semantics [24]. We extend these studies by probing predictions at all these levels simultaneously. This is important because it allows to control for correlations between levels – since words that are, for instance, syntactically surprising are, on average, also semantically surprising. Moreover, prior modelling of feature-specific predictions used domain-specific models that had to be independently trained, and typically incorporated linguistic context in a limited way. By contrast, our method (Figure 3) allows to derive multiple predictions from a single, large pre-trained model (like GPT-2) which has a much deeper grasp of linguistic context. However, a limitation of this method is that the resulting predictions are not independent. Therefore, you cannot test if levels interact without *also* creating a separate, domain-specific model. As such, the disentangling approach we used is complementary to the domain-specific modelling approach. Future work could combine the two, for instance to test if the hierarchical prediction we observed for phonemes applies to all linguistic levels – or whether predictions at some levels (e.g. syntax) might be independent.

In this study, we combined group-level analysis (of the EEG data) and individual-level analysis (of the MEG data). These approaches are complementary. While including more participants allows one to perform population-level inference, acquiring more data per participant allows one to evaluate effects within individuals. By combining both forms of analysis, we found that on the one hand, the basic effects of prediction and the comparison of hypotheses about its computational nature (probabilistic prediction, hierarchical prediction) were identical within and across each individual (Figure 2, 6, S5). But on the other hand, the exact spatiotemporal characteristics of these effects showed substantial variability (Figure 4, 5, S4, S7-S14). This suggest that while the prediction effects themselves at the EEG group-level are likely present in each individual, the precise spatiotemporal signatures (Figure 5) are probably best understood as a statistical average that is not necessarily representative of underlying individuals.

Because our analysis focused on evoked responses, we chose to probe predictions indirectly: via the neural markers of deviations from these predictions. As such, we cannot rule out that the effects might partly reflect ‘postdiction’. However, a purely postdictive explanation appears unlikely as it implies that after recognition, the brain computes a prediction of the recognised stimulus based on information available *before* recognition. While the data therefore indirectly support pre-activation, the representational format of these pre-activations is still an open question. In our analyses – and many theoretical models [6, 49]) – predictions are formalised as *explicit* probability distributions, but this is almost certainly a simplification. It remains unclear whether the brain represents probabilities implicitly. Alternatively, it might use a kind of approximation: graded, anticipatory processing that is perhaps functionally equivalent to probabilistic processing, but avoids having to represent (and compute with) probabilities. A potential way to address this question is to try to decode predictions before word onset [58]. Interestingly, this approach could be extended to assess whether predicted probabilities are represented before onset at different levels of the linguistic hierarchy, to test whether and which predicted distributions are reflected in pre-stimulus activity.

Why would the brain constantly predict upcoming language? Three – mutually non-exclusive – functions have been proposed. First, predictions can be used for *compression*: if predictable stimuli are represented succinctly, this yields an efficient code [6, 49] – conversely, optimising efficiency can make predictive coding emerge in neural networks [59]. A second, perhaps more studied function is that predictions can guide *inference*. Our analysis only probed prediction errors, and hence does not speak directly to such inferential effects of prediction – but earlier work suggests that linguistic context can indeed enhance neural representations in a top-down fashion [60, 61]; but see [62, 63]. Finally, predictions may guide *learning*: prediction errors can be used to perform error-driven learning without supervision. While learning is perhaps the least-studied function of linguistic prediction in cognitive neuroscience (but see [16]), it is its primary application in Artificial Intelligence [64, 65]. In fact, the language model we used (GPT-2) was created to study such predictive learning. These models are trained only to predict words, but learn about language more broadly, and can then be applied to practically any linguistic task [34, 65]. Interestingly, models trained with this predictive objective also develop representations that are ‘brain-like’, in the sense that they are currently the best encoders of linguistic stimuli to predict brain responses [66–69]. And yet, these predictive models are also brain-unlike in an interesting way – they predict upcoming language only at a single (typically lexical) level.

When prediction is used for compression or inference, it seems useful to predict at multiple levels, since redundancies and ambiguities also occur at multiple levels. But if predictions drive learning, why would the brain predict at multiple levels, when effective learning can be achieved using simple, single-level prediction? One fascinating option is that it might reflect the brain’s way to perform credit assignment within biological constraints. In artificial networks, credit assignment is typically done by first *externally* computing a single, global error term, and then ‘backpropagating’ this error through all levels of the network – but both these steps are biologically implausible [70]. Interestingly, it has been shown that hierarchical predictive coding networks can approximate or even implement classical back-propagation while using only Hebbian plasticity and local error computation [6, 70, 71]. Therefore, if the brain uses predictive error-driven learning, one might expect such prediction to be hierarchical, so error-terms can be locally computed throughout the hierarchy – which is in line with what we find.

Beyond the domain of language, there have been other reports of hierarchies of neural prediction, but these have been limited to artificial, predictive tasks or to restricted representational spans, such as successive stages in the visual system [72–74]. Our results demonstrate that even during passive listening of natural stimuli, the brain is engaged in prediction across disparate levels of abstraction (from speech sounds to meaning) based on timescales separated by three orders of magnitude (hundreds of milliseconds to minutes). These findings provide important evidence for hierarchical predictive processing in cortex. As such, they highlight how language processing in the brain is shaped by a domain-general neurocomputational principle: the prediction of perceptual inputs across multiple levels of abstraction.

## Methods

We analysed EEG and source localised MEG data from two experiments. The EEG data is part of a public dataset that has been published about before [27].

### Participants

All participants were native English speakers. In the EEG experiment, 19 subjects (13 male) between 19 and 38 years old participated; in the MEG experiment, 3 subjects participated (2 male) aged 35, 30, and 28. Both experiments were approved by local ethics committees (EEG: ethics committee of the School of Psychology at Trinity College Dublin; MEG: CMO region Arnhem-Nijmegen).

### Stimuli and procedure

In both experiments, participants were presented continuous segments of narrative speech extracted from audiobooks. The EEG experiment used a recording of Hemingway’s *The Old Man and the Sea*. The MEG experiment used 10 stories from the *The Adventures of Sherlock Holmes* by Arthur Conan Doyle. In total, EEG subjects listened to ∼1 hour of speech (containing ∼11,000 words and ∼35,000 phonemes); MEG subjects listened to ∼9 hours of speech (containing ∼85,000 words and ∼290,000 phonemes).

In the EEG experiment, each participants performed only a single session, which consisted of 20 runs of 180s long, amounting to the first hour of the book. Participants were instructed to maintain fixation and minimise movements but were otherwise not engaged in any task.

In the MEG experiment, each participant performed a total of ten sessions, each ∼1 hour long. Each session was subdivided in 6-7 runs of roughly ten minutes, although the duration varied as breaks only occurred at meaningful moments (making sure, for example, that prominent narrative events were not split across runs). Unlike in the EEG experiment, participants in the MEG dataset participants were asked to listen attentively and had to answer questions in between runs: one multiple choice comprehension question, a question about story appreciation (scale 1-7) and a question about informativeness.

### MRI acquisition and headcast construction

To produce the headcast, we needed to obtain accurate images of the participants’s scalp surface, which were obtained using structural MRI scans with a 3T MAGNETOM Skyra MR scanner (Siemens AG). We used a fast low angle shot (FAST) sequence with the following image acquisition parameters: slice thickness of 1 mm; field-of-view of 256 × 256 × 208 mm along the phase, read, and partition directions respectively; TE/TR = 1.59/4.5 ms.

### Data acquisition and pre-processing

The EEG data were originally acquired using a 128-channel (plus two mastoid channels) using an ActiveTwo system (BioSemi) at a rate of 512 Hz, and downsampled to 128 Hz before being distributed as a public dataset. We visually inspected the raw data to identify bad channels, and performed independent component analysis (ICA) to identify and remove blinks; rejected channels were linearly interpolated with nearest neighbour interpolation using MNE-python.

The MEG data were acquired using a 275 axial gradiometer system at 1200 Hz. For the MEG data, preprocessing and source modelling was performed in MATLAB 2018b using fieldtrip [75]. We applied notch filtering (Butterworh IIR) at the bandwidth of 49–51, 99–101, and 149– 151 Hz to remove line noise. Artifacts related to muscle contraction and squidjumps were identified and removed using fieldtrip’s semi-automatic rejection procedure. The data were downsampled to 150 Hz. To identify and remove eye blink artifacts, ICA was performed using the FastICA algorithm.

For both MEG and EEG analyses, we focus on the slow, evoked response and hence restricted our analysis to low-frequency components. To this end, we filtered the data between 0.5 and 8 Hz using a bidirectional FIR bandpass filter. Restricting the analysis to such a limited range of low frequencies (which are known to best follow the stimulus) is common when using regression ERP or TRF analysis, especially when the regressors are sparse impulses [28,31, 36]. The particular upper bound of 8 Hz is arbitrary but was based on earlier papers using the same EEG dataset to study how EEG tracks acoustic and linguistic content of speech [31, 56, 61].

### Head and source models

The MEG sensors were co-registered to the subjects’ anatomical MRIs using position information of three localization coils attached to the headcasts. To create source models, FSL’s Brain Extraction Tool was used to strip non-brain tissue. Subject-specific cortical surfaces were reconstructed using Freesurfer, and post-processing (downsampling and surface-based alignment) of the reconstructed cortical surfaces was performed using the Connectome Workbench command-line tools (v 1.1.1). This resulted in cortically-constrained source models with 7,842 source locations per hemisphere. We created single-shell volume conduction models based on the inner surface of the skull to compute the forward projection matrices (leadfields).

### Beamformer and parcellation

To estimate the source time series from the MEG data, we used linearly constrained minimum variance (LCMV) beamforming, performed separately for each session, using Fieldtrip’s ft_sourceanalysis routine. To reduce the dimensionality, sources were parcellated, based on a refined version of the Conte69 atlas, which is based on Brodmann’s areas. We computed, for each session, parcel-based time series by taking the first principal component of the aggregated time series of the dipoles belonging to the same cortical parcel.

### Self-attentional language model

Contextual predictions were quantified using GPT-2 – a large, pre-trained language model [34]. Formally, a language model can be cast as a way of assigning a probability to a sequence of words (or other symbols), (*x*_1_, *x*_2_, …, *x*_*n*_). Because of the sequential nature of language, the joint probability, *P* (*X*) can, via the chain rule, be factorised as the product of conditional probabilities:

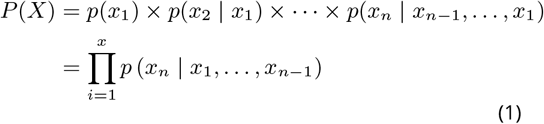

Since the advent of neural language models, as opposed to statistical (Markov) models, methods to compute these conditional probabilities have strongly improved. Improvements have been especially striking in the past two years with the introduction of the *Transformer* [76] architecture, which allows efficient training of very large networks on large, diverse data. This resulted in models that dramatically improved the state-of-the art in language modelling on a range of domains.

GPT-2 [34] is one of these large, transformer-based language models and is currently among the best publcicly released models of English. The architecture of GPT-2 is based on the decoder-only version of the transformer. In a single forward pass, it takes a sequence of tokens *U* = (*u*_1_, …, *u*_*k*_) and computes a sequence of conditional probabilities, (*p*(*u*_1_), *p*(*u*_2_|*u*_1_), …, *p*(*u*_*k*_ | *u*_1_, …, *u*_*k*−1_)). Roughly, the full model (see Figure S1) consists of three steps: first, an embedding step encodes the sequence of symbolic tokens as a sequence of vectors which can be seen as the first hidden state *h*_*o*_. Then, a stack of transformer blocks, repeated *n* times, each apply a series of operations resulting in a new set of hidden states *h*_*l*_, for each block *l*. Finally, a (log-)softmax layer is applied to compute (log-)probabilities over target tokens. Formally, then, the model can be summarised in three equations:

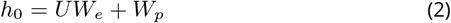

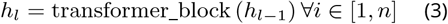

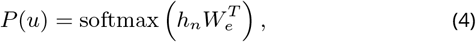

where *W*_*e*_ is the token embedding and *W*_*p*_ is the position embedding (see below).

The most important component of the transformer-block is the *masked multi-headed self-attention* (Fig S1). The key operation is self-attention, a seq2seq operation turning a sequence of input vectors (x_1_, x_2_, … x_*k*_) into a sequence of output vectors (y_1_, y_2_, …, y_*k*_). Fundamentally, each output vector y_*i*_ is a weighted average of the input vectors: 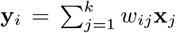. Critically, the weight *w*_*i,j*_ is not a parameter but is *derived* from a function over input vectors x_*i*_ and x_*j*_. The Transformer uses (scaled) *dot product attention*, meaning that the function is simply a dot product between the input vectors 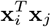, passed through a softmax make sure that the weights sum to one, scaled by a constant determined by the dimensionality, 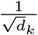 (to avoid the dot-products growing too large in magnitude): 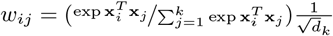.

In self-attention, then, each input x_*i*_ is used in three ways. First, it is multiplied by the other vectors to derive the weights for its own output, y_*i*_ (as the *query*). Second, it is multiplied by the other vectors to determine the weight for any other output y_*j*_ (as the *key*). Finally, to compute the actual outputs it is used in the weighted sum (as the *value*). Different (learned) linear transformations are applied to the vectors in each of these use cases, resulting in the Query, Key and Value matrices (*Q, K, V*). Putting this all together, we arrive at the following equation:

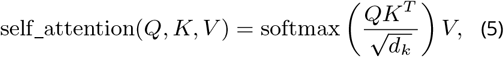

where *d*_*k*_ is dimension of the keys/queries. In other words, self_attention simply computes a weighted sum of the values, where the weight of each value is determined by the dot-product similarity of the query with its key. Because the queries, keys and values are linear transformations of the same vectors, the input *attends itself*.

To be used as a language model, two elements need to be added. First, the basic self-attention operation is not sensitive to the order of the vectors: if the order of the input vectors is permuted, the output vectors will be identical (but permuted). To make it position-sensitive, a position embedding *W*_*p*_ is simply added during the embedding step – see Equation 2. Second, to enforce that the model only uses information from one direction (i.e left), a mask is applied to the attention weights (before the softmax) which sets all elements above the diagonal to −∞. This makes the self-attention *masked*.

To give the model more flexibility, each transformer block actually contains multiple instances of the basic self-attention mechanisms from (5). Each instance (each *head*) applies different linear transformations to turn the same input vectors into a different set of *Q, K* and *V* matrices, returning a different set of output vectors. The outputs of all heads are concatenated and then reduced to the initial dimensionality with a linear transformation. This makes the self-attention *multi-headed*.

In total, GPT-2 (XL) contains *n* = 48 blocks, with 12 heads each; a dimensionality of *d* = 1600 and a context window of *k* = 1024, yielding a total 1.5×10^9^ parameters. We used the PyTorch implementation of GPT-2 provided by HuggingFace’s *Transformers* package [77].

### Lexical predictions

We passed the raw texts through GPT-2 (Equations 2-4) for each run independently (assuming that listeners’ expectations would to some extent ‘reset’ during the break). This resulted in a (log-)probability distribution over tokens *P* (*U*). Since GPT-2 uses Byte-Pair Encoding, a token can be either punctuation or a word or (for less frequent words) a word-part. How many words actually fit into a context window of length *k* therefore depends on the text. For words spanning multiple tokens, we computed word probabilities simply as the joint probability of the tokens. ‘For window-placement, we used the constraint that the windows had an overlap of at least 700 tokens, and that they could not start mid-sentence (ensuring that the first sentence of the window was always well-formed).

As such, for each word *w*_*i*_ we computed *p*(*w*_*i*_|context), where ‘context’ consisted either of all preceding words in the run, or of a sequence of prior words constituting a well-formed context that was at least 700 tokens long.

### Syntactic and semantic predictions

Feature-specific predictions were computed from the lexical prediction. To this end, we first truncated the unreliable tail from the distribution using a combination of top-k and nucleus truncation. The nucleus was defined as the “top” *k* tokens with the highest predicted probablility, where *k* was set dynamically such that the cumulative probability was at least 0.9. To have enough information also for very low entropy cases (where *k* becomes small), we forced *k* to be a least 40.

From this truncated distribution, we derived feature-specific predictions by analysing the predicted words. For the syntactic predictions, we performed part of speech tagging on every potential sentence (i.e. the context plus the predicted word) with Spacy to derive the probability distribution over parts-of-speech, from which the syntactic surprise was calculated as the negative log probability of the POS of a word, − log(*P* (POS_*n*_ |context)).

For the semantic prediction, we took a weighted average of the glove embeddings of the predicted words to compute the expected vector: 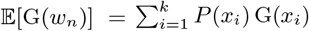, where G(*w*_*i*_) is the GloVe embedding for predicted word *w*_*i*_. From this prediction, we computed the semantic prediction error as the cosine distance between the predicted and observed vector:

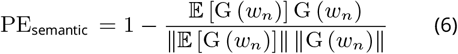

### Phonemic predictions

Phonemic predictions were formalised in the context of incremental word recognition [27, 29]. This process can be cast as probabilistic prediction by assuming that brain is tracking the *cohort* of candidate words consistent with the phonemes so far, each word weighted by its prior probability. We compared two such models that differed only in the prior probability assigned to each word.

The first model was the single-level or frequency-weighted model (Fig 6), in which prior probability of words was fixed and defined by a word’s overall probability of occurrence (i.e. lexical frequency). The probability of a specific phoneme (*A*), given the prior phonemes within a word, was then calculated using the statistical definition:

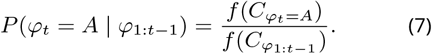

Here, *f* (*C*_*ϕ*__*t*=*A*_) denotes the cumulative frequency of all words in the remaining cohort of candidate words if the next phoneme were *A*, and 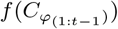denotes the cumulative frequency of all words in the prior cohort (equivalent to *f* (*C*) of all potential continuations). If a certain continuation did not exist and the cohort was empty, *f* (*C*_*ϕt*=*A*_) was assigned a laplacian pseudocount of 1. To efficiently compute (7) for every phoneme, we constructed a statistical phonetic dictionary as a digital tree that combined frequency information from SUBTLEX database and pronunciation from the CMU dictionary.

The second model was equivalent to the first model, except that the prior probability of each word was not defined by its overall probability of occurrence, but by its conditional probability in that context (based on GPT-2). This was implemented by constructing a separate phonetic dictionary for every word, in which lexical frequencies were replaced by implied counts derived from the lexical prediction. We truncated the unreliable tail from the distribution and replaced that by a flat tail that assigned each word a pseudocount of 1. This greatly simplifies the problem as it only requires to assign implied counts for the top *k* predicted words in the dynamic nucleus. Since all counts in the tail are 1, the cumulative implied counts of the nucleus is complementary to the the length of the tail, which is simply the difference between the vocabulary size and nucleus size (*V* − *k*). As such a little algebra reveals:

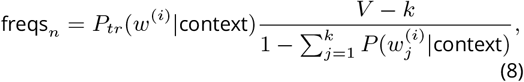

where *P*_*tr*_(*w*^(*i*)^ |context) is the trunctated lexical lexical prediction, and 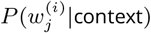 is predicted probability that word *i* in the text is word *j* in the sorted vocabulary.

Although we computed probabilities using the simple statistical definition of probability, these two ways of assigning lexical frequencies are equivalent to two kinds of priors in a Bayesian model. Specifically, in the first model the prior over words is the fixed unconditional word probability, while in the second model the prior is the contextual probability, itself based on a higher level (lexical) prediction. This makes the second computation *hierarchical* because phoneme predictions are based on not just (at the first level) on short sequences of within-word phonemes, but also on a contextual prior which itself (at the second level) is based on long sequences of prior words.

### Non-predictive control variables

To ensure we were probing effects of predictions, we had to control for various non-predictive variables: onsets, acoustics, frequency and semantic congruency. We will briefly outline our definitions of each.

For speech, it is known that the cortical responses are sensitive to fluctuations in the envelope – which is specifically driven by rapid increases of the envelope amplitude (or ‘acoustic edges’) [78]. To capture these fluctuations in a sparse, impulse-based regressor we quantified the amplitude of these edges as the variance of the envelope over each event (e.g. phoneme) following [61]. A second non-predictive variable is frequency. We accounted for frequency as the overall base rate or unconditional probability of a word, defining it similarly to lexical surprise as the unigrams surprise − log *P*(word) based on its frequency of occurrence in subtlex.

The final non-predictive variable was semantic congruency or integration difficulty. This speaks to the debate wether effects of predictability reflect prediction or rather post-hoc effects arising when integrating a word into the semantic context. This can be illustrated by considering a constraining context (‘coffee with milk and … ‘). When we contrast a highly expected word (‘sugar’) and an unexpected word (e.g. ‘dog’), the unexpected word is not just less likely, but also semantically incongruous in the prior context. As such, the increased processing cost reflected by effects like N400 increases might not (only) be due to a violated *prediction* but due to difficulty integrating the target word (‘dog’) in the semantic context (‘coffee with milk’) [7, 18, 51, 52]. As a proxy for semantic integration difficulty we computed the semantic congruency of a word in its context defined as the cosine dissimilarity (see (6)) between the average semantic vector of the prior context words and the target content word, following [31]. This metric is known to predict N400-like modulations and can hence capture the extent to which such effects can be explained by semantic congruency only [31, 52].

### Word-level regression models

The word-level models (see Fig S2 for graphical representation) captured neural responses to words as a function of word-level variables. The *baseline* model formalised the hypothesis that responses to words were not affected by word unexpectedness but only by the following non-predictive confounds: word onsets, envelope variability (acoustic edges), semantic congruency (integration difficulty) and word frequency.

The *probabilistic prediction* model formalised the hypothesis that predictions were continuous and probabilistic. This model was identical to the baseline model plus the lexical surprise (or negative log probability of a word), for every word. This was based on normative theories of predictive processing which state that the brain response to a stimulus should be proportional to the negative log probability of that stimulus [6].

The *constrained guessing* model formalised the classical psycholinguistic notion of prediction as the all-or-none preactivation of specific words in specific (highly constraining) contexts [33]. We translated the idea of all-or-none prediction into a regression model using an insight by Smith and Levy which implied that all-or-none predictions result in a linear relationship between word probability and brain responses [9]. The argument follows from two assumptions: (1) all predictions are all-or-none; and (2) incorrect predictions incur a cost, expressed as a prediction error brain response (fixed in size because of assumption 1). For simplicity, we first consider the unconstrained case (i.e. subjects make a prediction for *every* stimulus), and we bracket all other factors affecting brain responses by absolving them into an average brain response, *y*_baseline_. As such, the response to any word is either *y*_baseline_ (if the prediction is correct) or *y*_baseline_ + *y*_error_ (if it was false). For any individual stimulus, this equation cannot be used (as we don’t know what a subject predicted). But if we assume that predictions are approximately correct, then the probability of a given prediction to be incorrect simply becomes ∼(1 − *p*). As such, *on average*, the response becomes *y*_resp_ = *y*_baseline_ + (1 − *p*)*y*_error_. In other words, a linear function of word improbability. To extend this to the constrained case, we only define the improbability regressor for constraining contexts, and add a constant to those events to capture (e.g. suppressive) effects of correct predictions (Figure S2). To identify ‘constraining contexts’, we simply took the 10% of words with the lowest prior lexical entropy. The choice of 10% was arbitrary – however, using a slightly more or less stringent definition would not have changed the results because the naive guessing model (which included linear improbability for *every* word) performed so much better (see Figure S5).

### Integrated regression model

For all analyses on feature-specific predictions, we formulated an integrated regression model with both word-level and phoneme-level regressors (Figure S6). To avoid collinearity between word and phoneme level regressors, phoneme-level regressors were only defined for word-non-initial phonemes, and word-level regressors were define for word-onset. As regressors of interest this model included phonemic surprise, syntactic surprise and semantic prediction error. In principle, we could have also included phoneme and syntactic entropy rather than just surprise (e.g. [13]) – however, these were highly correlated with the respective surprise. Since this was already a complex regression model, including more correlated regressors would have made the coefficients estimates less reliable and hence more difficult to interpret. As such, we did not include both but focussed on surprise because it has the most direct relation to stimulus evoked effect.

### Phoneme-level regression models

To compare different accounts of phoneme prediction, we formulated three regression models with only regressors at the individual phoneme level (Figure S15). In all models, following [27] we used separate regressors for word-initial and word-non-initial phonemes, to account for juncture phonemes being processed differently. The baseline model only included non-predictive factors of word-boundaries, phoneme onsets, envelope variability, and uniqueness points. The two additional models also included phoneme surprise and phoneme entropy from either the hierarchical model or non-hierarchical model. To maximise our ability to dissociate the hierarchical prediction and non-hierarchical prediction, we included both entropy and surprise. Although these metrics are correlated, adding both should add more information to the model-comparison, assuming that there is some effect of entropy [13]. (Note that here, we were only interested in model comparison, and not in comparing the coefficients, which may become more difficult when including both.)

### Time resolved regression

As we were interested in the evoked responses, variables were regressed against EEG data using time-resolved regression, within a regression ERP/F (or impulse TRF) framework [31, 35]. Briefly, this involves using impulse regressors for both constants and covariates defined at event onsets, and then temporally expanding the design matrix such that each predictor column *C* becomes a series of columns over a range of temporal lags 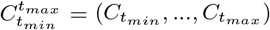. For each predictor one thus estimates a series of weights 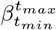 (Fig 1) which can be understood as the *modulation function* describing how a given regressor modulates the neural response over time, and which corresponds to the *effective* evoked response that would have been obtained in a time-locked ERP/ERF design. Here, we used a range between -0.2 and 1.2 seconds. All data and regressors were standardised and coefficients were estimated with *𝓁*_2_-norm regularised (Ridge) regression:

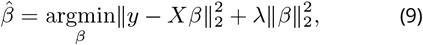

using the scikit learn sparse matrix implementation. In both datasets, models were estimated by concatenating the (time-expanded) design matrix across all runs and sessions. Regularisation was set based on leave-one-run-out *R*^2^ comparison; for inference on the weights in the EEG data this was done across subjects to avoid doing statistics over coefficients with different amounts of shrinkage.

### Model comparison

In both datasets, model comparison was based on comparing cross-validated correlation coefficients. Cross-validation was performed in a leave-one-run-out cross-validation scheme, amounting to 19-fold cross-validation in the EEG data and between 63 and 65-fold cross-validation for the MEG data (in some subjects, some runs were discarded due to technical problems).

For the EEG data, models’ cross-validated prediction performance was performed across subjects to perform population-level inference. To this end, we reduced the scores into a single *n*_*subs*_ dimensional vector by taking the median across folds and the mean across channels. Critically, we did not select any channels but used the average across the scalp. For the MEG data, models were only statistically compared on a within within-subject basis. Because the MEG data was source localised we could discard sources known to be of no interest (e.g. early visual cortex). To this end, we focussed on the language network, using a rather unconstrained definition encompassing all Brodmann areas in the temporal lobe, plus the temporo-parietal junction, and inferior frontal gyrus and dorsolateral prefrontal cortex; all bilaterally (see Figure S16).

### Statistical testing

All statistical tests were two-tailed and used an alpha of 0.05. For all simple univariate tests performed to compare model-performance within and between subjects, we first verified that the distribution of the data did not violate normality and was outlier free, determined by the D’Agostino and Pearson’s test implemented in SciPy and the 1.5 IQR criterion, respectively. If both criteria were met, we used a parametric test (e.g. paired t-test); otherwise, we resorted to a non-parametric alternative (e.g. Wilcoxon sign rank).

In EEG, we performed mass-univariate tests on the coefficients across participants between 0 and 1.2 seconds. This was firstly done using cluster-based permutation tests [79, 80] to identify clustered significant effects as in Figure 5 (10,000 permutations per test). Because the clustered effects as in Figure 5 only provide a partial view, we also reported more comprehensive picture of the coefficients across all channels (Figure, S8 S3); there, we also provide multiple-comparison corrected p-values to indicate statistical consistency of the effects; these were computed using TFCE. In the MEG, multiple comparison correction for comparison of explained variance across cortical areas was done using Treshold Free Cluster Enhancement (TFCE). In both datasets, mass-univariate testing was performed based on one-sample t-tests plus the ‘hat’ variance adjustment method with *σ* = 10^−3^.

### Polarity-alignment

In the source localised MEG data, the coefficients in individuals (e.g. Figure S14 S11) are symmetric in polarity, with the different sources in a single response having an arbitrary sign due to ambiguity of the source polarity. To harmonise the polarities, and avoid cancellation when visualising the average coefficient, we performed a polarity-alignment procedure. This was based on first performing SVD, **A** = **AΣV**^⊤^, where **A** is the *m* × *n* coefficient matrix, with *m* being the number of sources and *n* the number of regressors; and then multiplying each row of **A** by the sign of the first right singular vector. Because the right singular vectors (columns of **U**) can be interpreted as the eigen vectors of the source-by-source correlation matrix, this can be thought of as flipping the sign of each source as a function of its polarity with respect to the dominant correlation. This procedure was used for visualisation purposes only (see Fig S4 and S11-S14).

## Data and code availability

Data and code to reproduce all results will be made public at the Donders Repository. The full MEG dataset will be made public in a separate resource publication.

## Acknowledgements

This work was supported by The Netherlands Organisation for Scientific Research (NWO Research Talent grant to M.H.; NWO Vidi grant to F.P.d.L.; NWO Vidi 864.14.011 to JMS; Gravitation Program Grant Language in Interaction no. 024.001.006 to P.H.) and the European Union Horizon 2020 Program (ERC Starting Grant 678286, ‘Contextvision’ to F.P.d.L). We wish to thank Michael P Broderick, Giovanni M. Di Liberto, and colleagues from the Lalor lab for making the EEG dataset openly available. We thank all the authors of the open source software we used and apologise for citation limits that prevent us from citing all tools used.

## Contributions

Conceptualisation: MH, FPdL, PH; Formal analysis: MH; Data collection: KA, JMS; Source modelling: KA, JMS; Original draft: MH; Final manuscript: MH, FPdL, PH, JMS, KA.

## Supplementary materials

**Figure S1.**
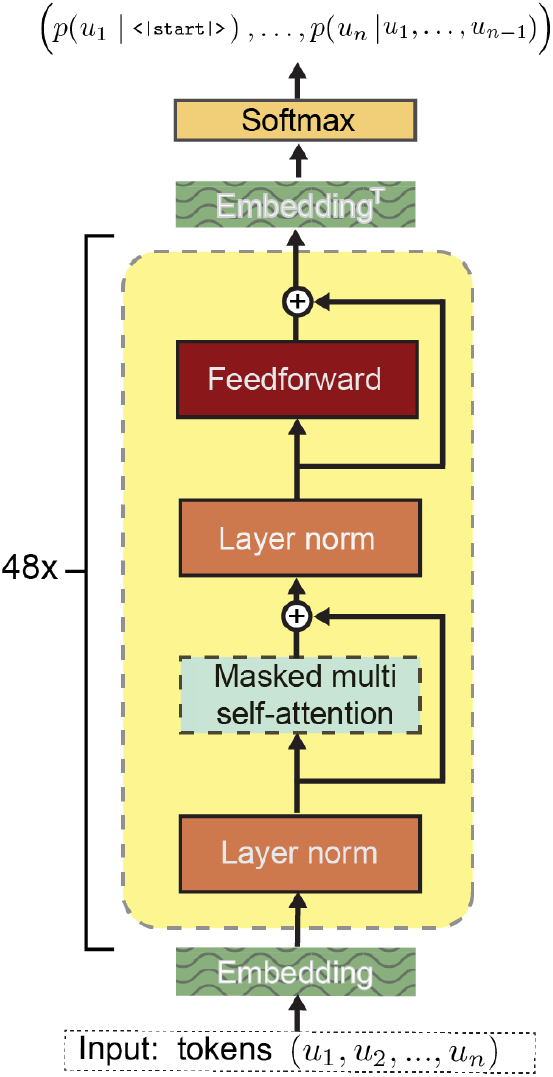
GPT-2 Architecture. Note that this panel is a re-rendered version of the original GPT schematic, slightly modifyied and re-arranged to match the architecture of GPT-2. For more details on the overall architecture and on the critical operation of self-attention, see *Methods*. In this graphic, Layer Norm refers to layer normalisation as described by Ba et al. Not visualised here is the initial tokenisation, mapping a sequence of characters into tokens.

**Figure S2.**
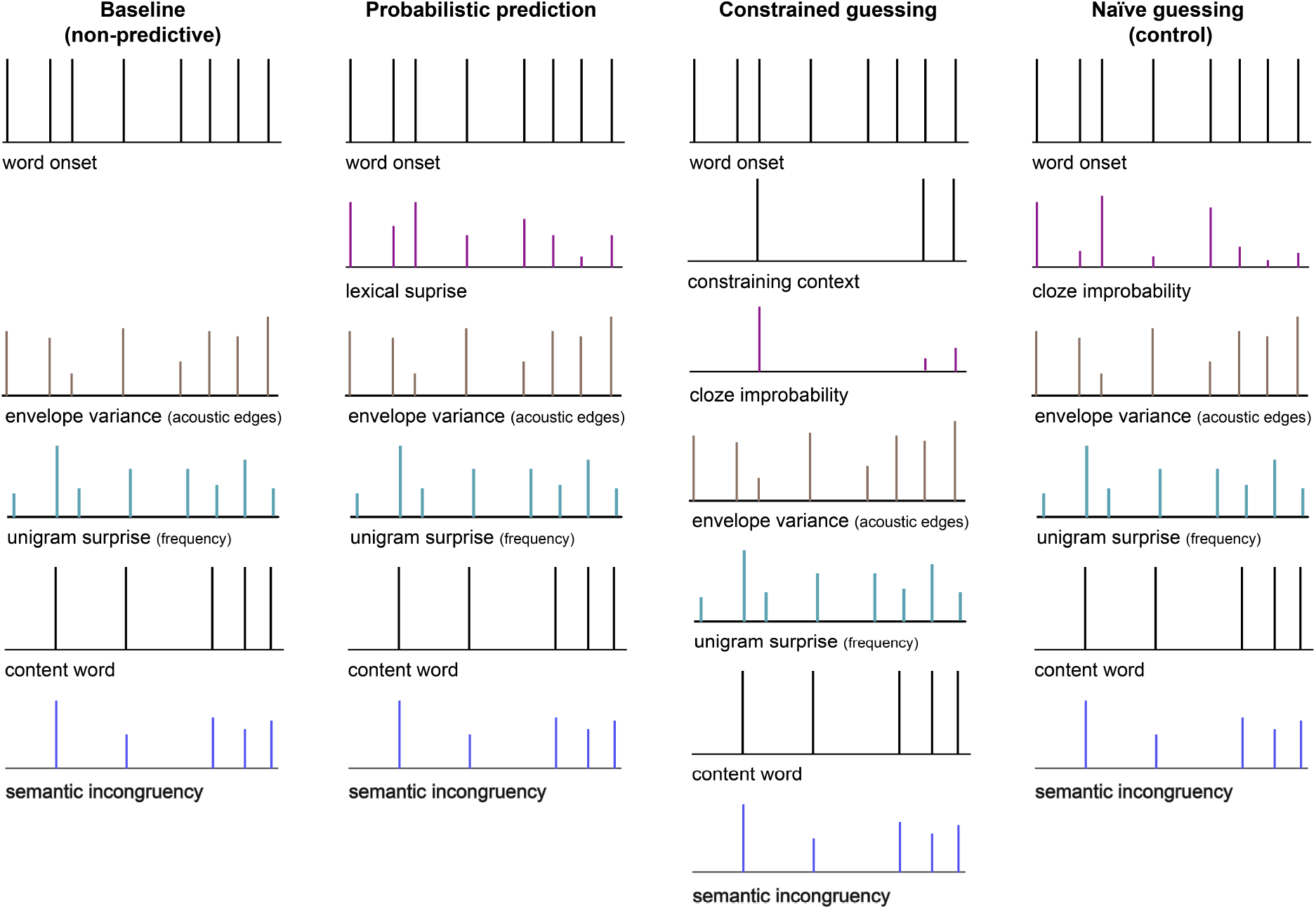
Word-level regression models. Schematic of the main models plus the control model of the initial model comparison to test for predictive processing at the word level. Because we use a regression ERP/ERF scheme [35], aimed at capturing (modulations of) the evoked response to discrete events like words or phonemes, all regressors are modelled as impulses (see *Methods*).

**Figure S3.**
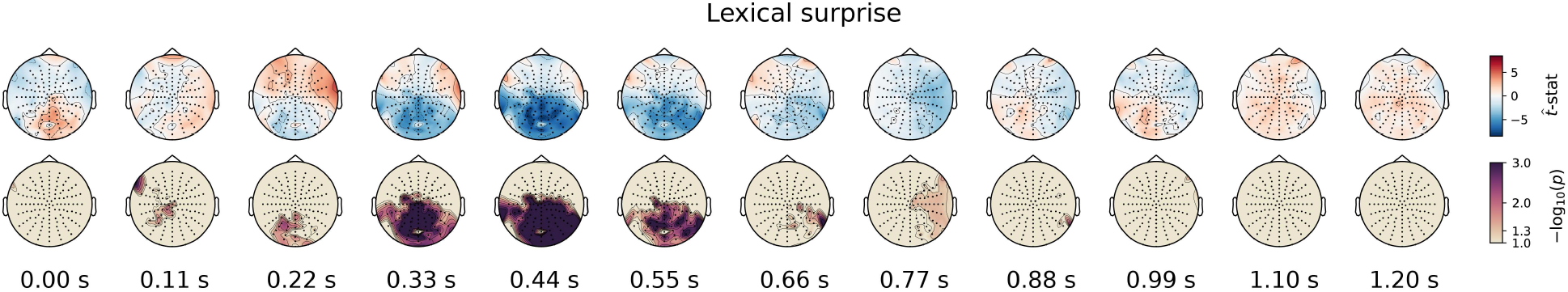
Full EEG topographies of the effects of lexical surprise. These topographies show the average t-statistics of the coefficients (upper row) and respective FWE-corrected significance (lower row) of the lexical surprise regressor from the *probabilistic prediction* model (Figure S2). As such, while Figure 2b shows the coefficients averaged over channels participating in the cluster (thereby only visualising *the effect*) these topographies visualise the results comprehensively across all channels over time.

**Figure S4.**
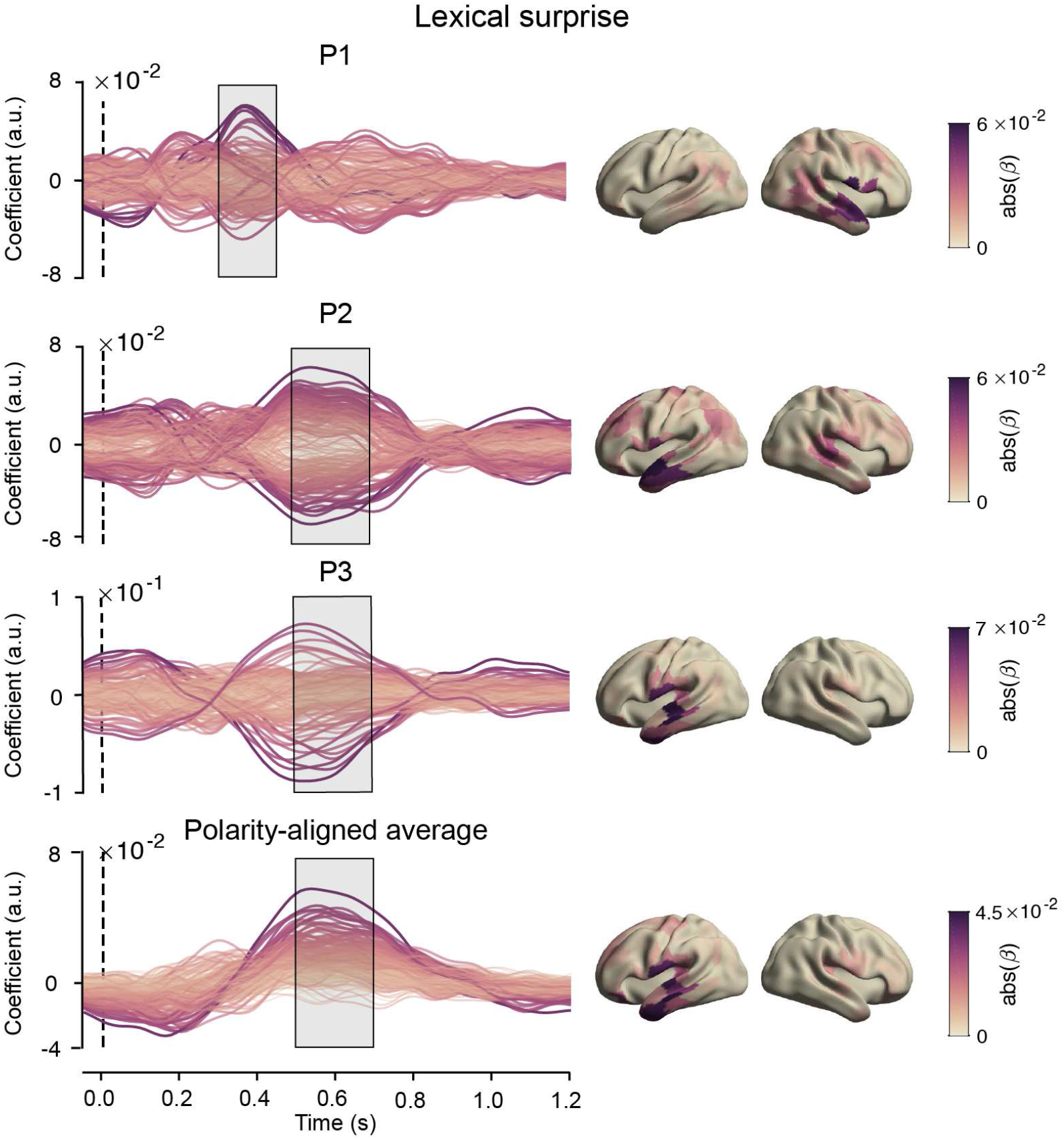
Coefficients for lexical surprise from the lexical model. (Figure S2) Left column: timecourses of the coefficients at each MEG source-localised parcel for lexical surprise for all MEG participants, and the polarity-aligned average across them. Right column: Absolute value of the coefficients averaged across the highlighted period plotted across the brain.

**Figure S5.**
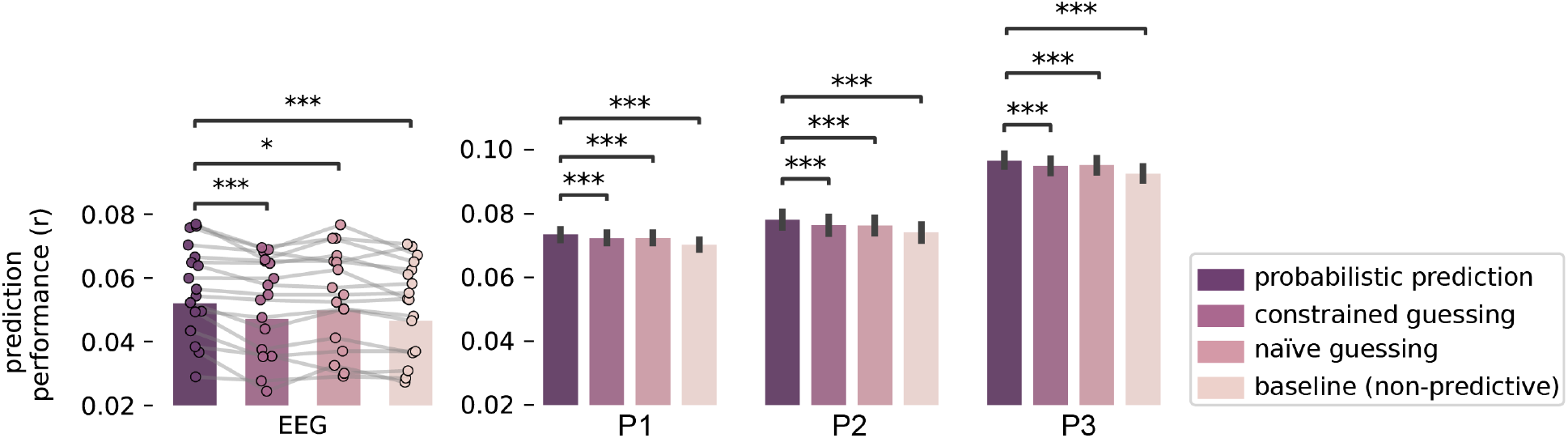
Model comparison results across all channels (EEG) and the full language network (MEG). Same as in Figure 2a, but now including the ‘naive guessing’ control model. Like the constrained guessing model, this model included a linear estimate of word probability, but defined for every word rather than only for constraining contexts. This model was introduced to identify which of the two differences between the *probabilistic prediction* and *constrained guessing* model. i.e. assuming that predictions are (i) categorical vs. probabilistic and (ii) occasional vs. continuous. made the largest difference in model performance. As can be seen, the *naive guessing* model performed considerably better than the *constrained guessing* model, but consistently worse than the *probabilistic prediction* model. This clearly shows that the modulatory effect of unexpectedness is not limited to only highly constraining contexts, but that that it applies much more generally – in line with the notion of continuous prediction. Strictly speaking, the naive guessing model formalises the hypothesis that the brain ‘naively’ makes *all-or-none* guesses about *every* upcoming word. Given that this hypothesis is a-priori so implausible, it may seem surprising that the model still performs comparably well. However, we should note that the probabilistic prediction regressor (*surprise*) and the categorical prediction regressor (linear (im)probability) are highly correlated (∼0.7) because one is a monotonic function of the other. Therefore, we suggest the results are better interpreted the other way around: the fact that. despite being so correlated. the log-probability is consistently a better linear predictor of neural responses than the linear probability clearly supports predictive processing theories, which postulate that the neural response to a stimulus should be proportional to negative log-probability of that stimulus.

**Figure S6.**
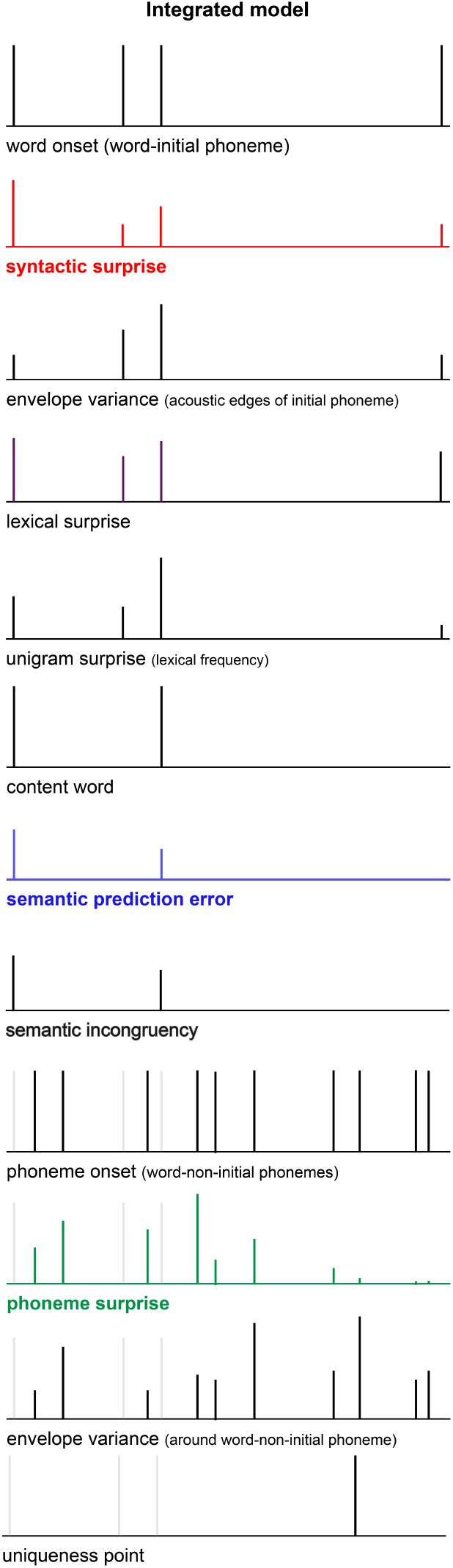
Regressors of the integrated feature-specific model. Same as Figure S5, but for the integrated feature-specific regression model. The three regressors of interest. syntactic surprise, semantic prediction error and phonemic surprise. are coloured, all control regressors are in black. Following the regression ERP/ERF scheme [35], aimed at capturing (modulations of) the evoked response to discrete events like words or phonemes, all regressors are modelled as impulses (see *Methods*). To avoid collinearity between word an and phoneme regressors, phoneme regressors (both events and covariates) are restricted to all non-initial phonemes.

**Figure S7.**
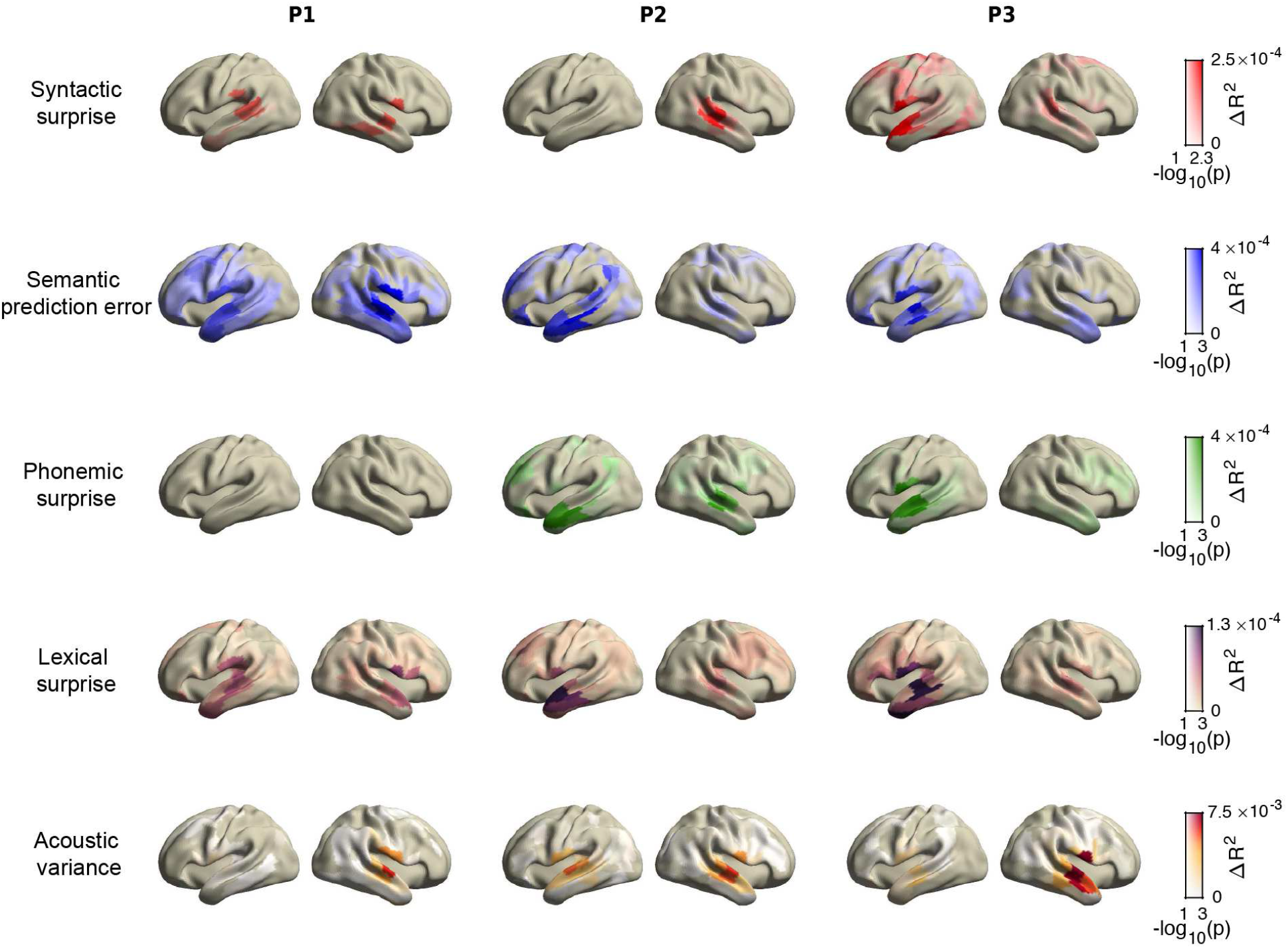
Unique explained variance for five regressors across the brain. Same as Figure 4, but including 2 control regressors (lexical surprise and acoustic variance) for comparison. Colours indicate amount of additional variance explained by each regressor; opacity indicates the FWE-corrected statitsical significance (across cross-validation folds). Note that *p <* 0.05 is equivalent to − log_10_(*p*) *>* 1.3.

**Figure S8.**
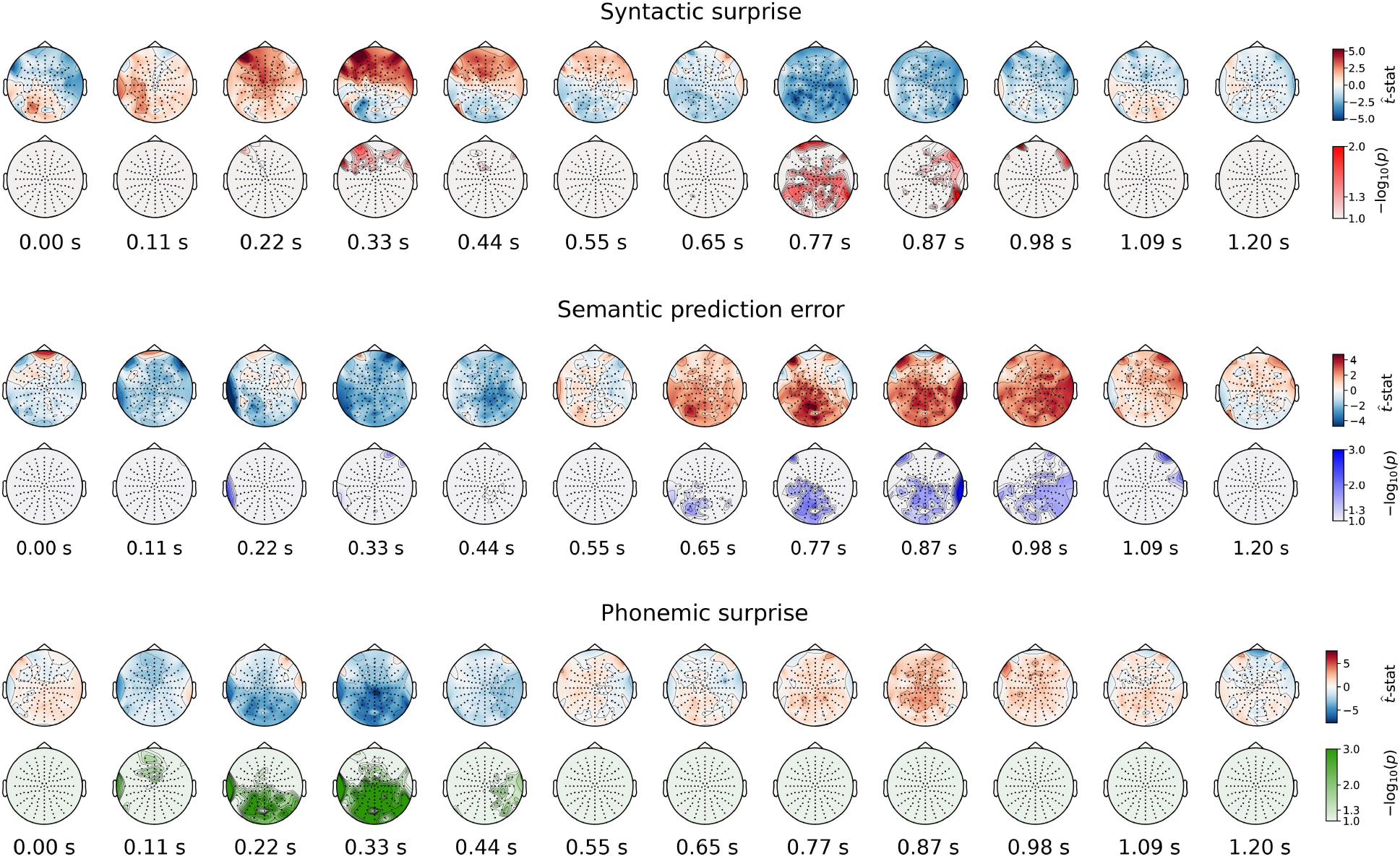
Full topographies of the coefficients and significance of feature-specific prediction errors. For each feature-specific prediction error regressor, the topographies show the t-statistics of the coefficients (upper row) and the respective TFCE-corrected significance (lower row). So while Figure 5 only shows the coefficients averaged over channels participating in the cluster (thereby only visualising *the effect*) these topographies visualise the results comprehensively across all channels, over time.

**Figure S9.**
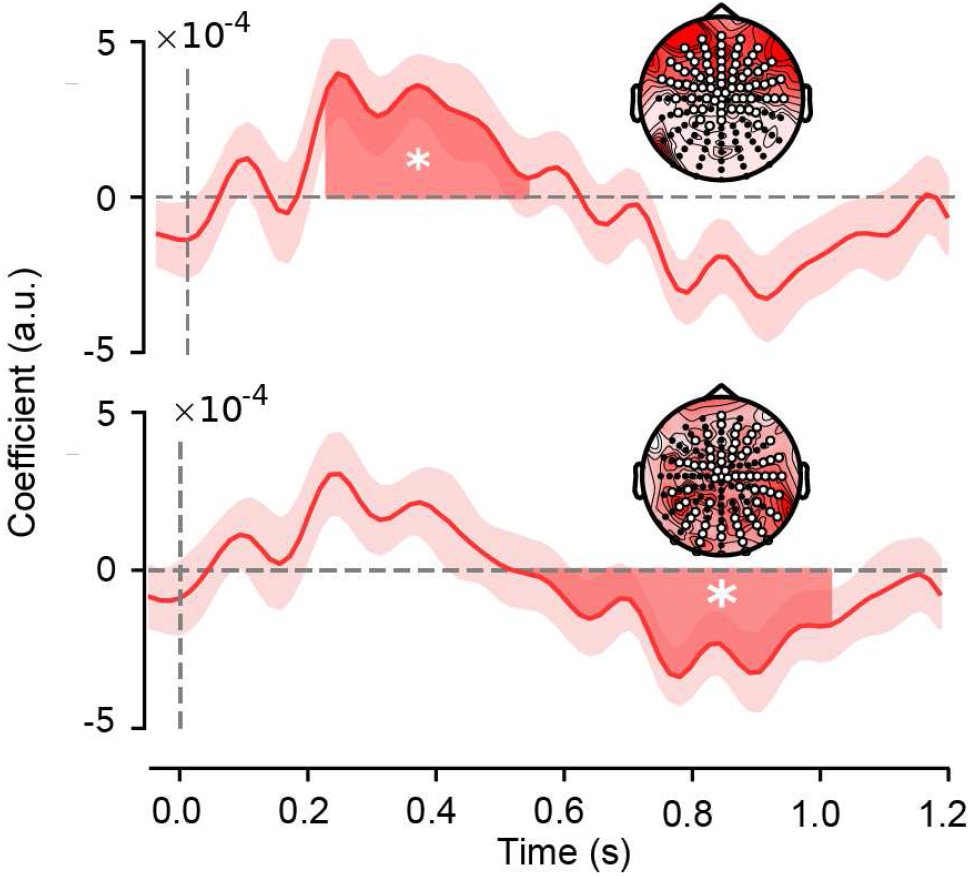
Significant effects of syntactic surprise in the EEG data. Two significant effects were observed in the modulation functions for syntactic surprise: an early positive effect with a frontal topography (upper panel) and a later negative effect based on a distributed cluster (lower panel). The early effect tightly replicates recent model-based studies on EEG effects of syntactic surprise, and was also found in the MEG data. By contrast, the late effect of syntactic surprise is not in line with any earlier study (note that it is negative unlike the syntactic P600) and importantly was not replicated in the MEG data. Therefore we only consider the early effect a ‘main’ effect of syntactic surprise (visualised in the main Figure 5) and we advice to refrain from interpreting the late effect before it is independently replicated.

**Figure S10.**
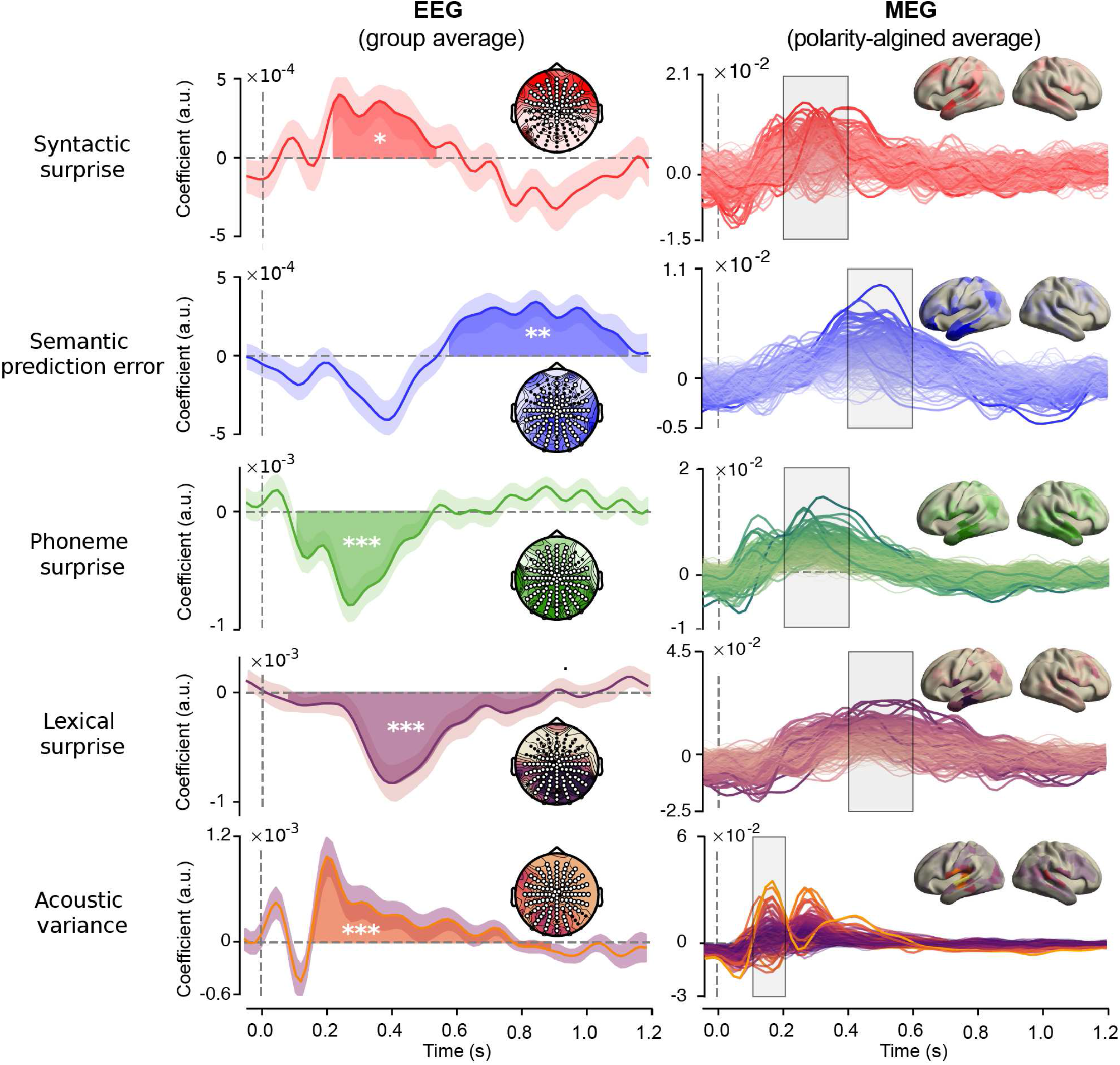
Coefficients for each prediction error, plus two control variables. EEG (left column): coefficient modulation function averaged across the channels participating for at least one sample in the significant clusters. Highlighted area indicates temporal extent of the cluster. Shaded area around waveform indicates bootstrapped standard errors. Stars indicate cluster-level significance; *p <* 0.05 (*), *p <* 0.05 (**), *p <* 0.001 (***). Insets represent channels assigned to the cluster (white dots) and the distribution of absolute values of t-statistics. MEG (right column): polarity aligned responses averaged across participants for all sources (same as in Figure 5 but without averaging over sources, and including two control variables). Insets represent topography of absolute value of coefficients averaged across the highlighted period. Note that due to polarity alignment, sign information is to be ignored for the MEG plots.

**Figure S11.**
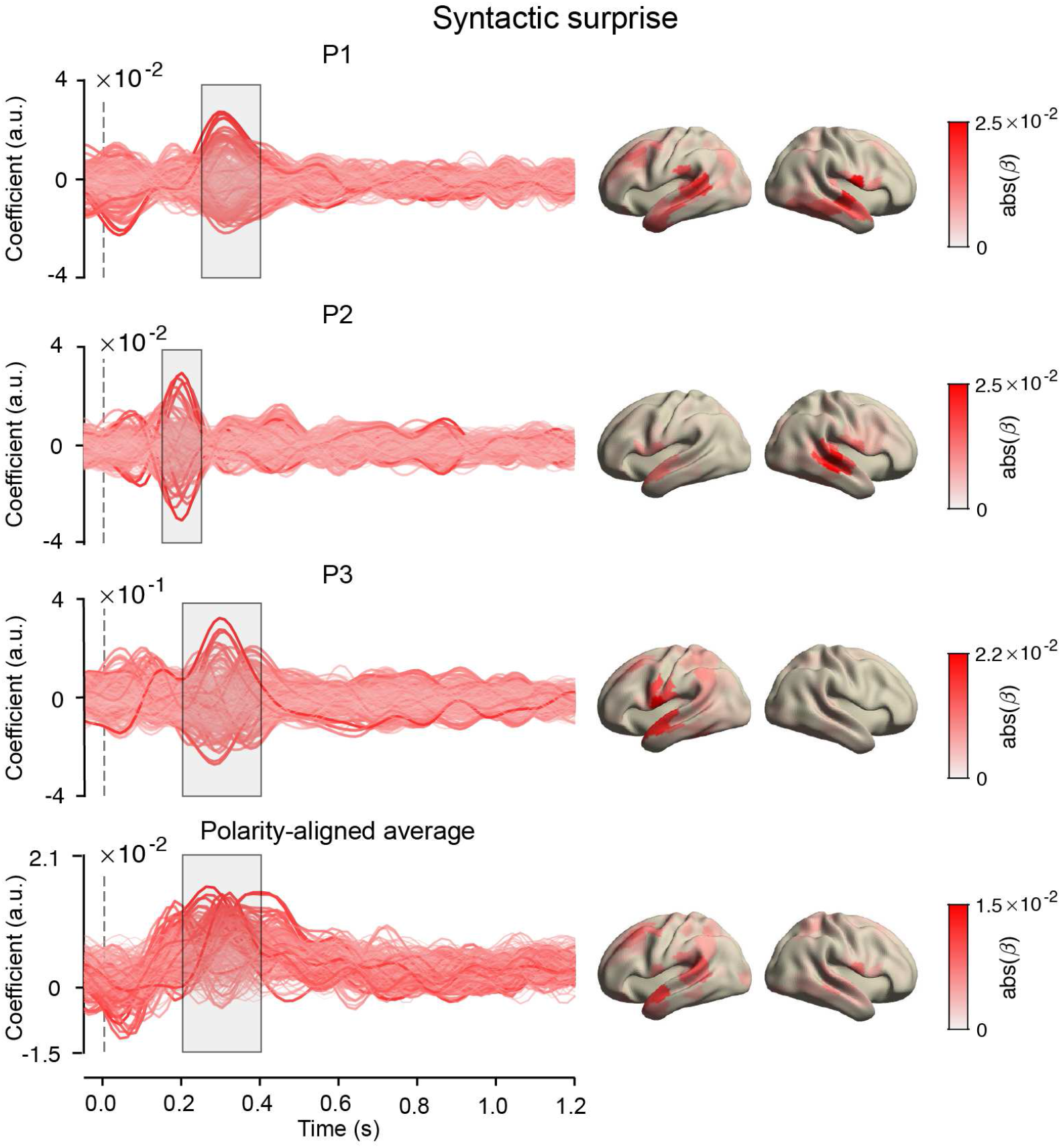
Coefficients for syntactic surprise from the integrated model (Figure S6) Left column: coefficients for each source for each individual in the MEG experiment, and the polarity-aligned average across participants. Right column: absolute value of the coefficients across the brain, averaged across the highlighted time-period.

**Figure S12.**
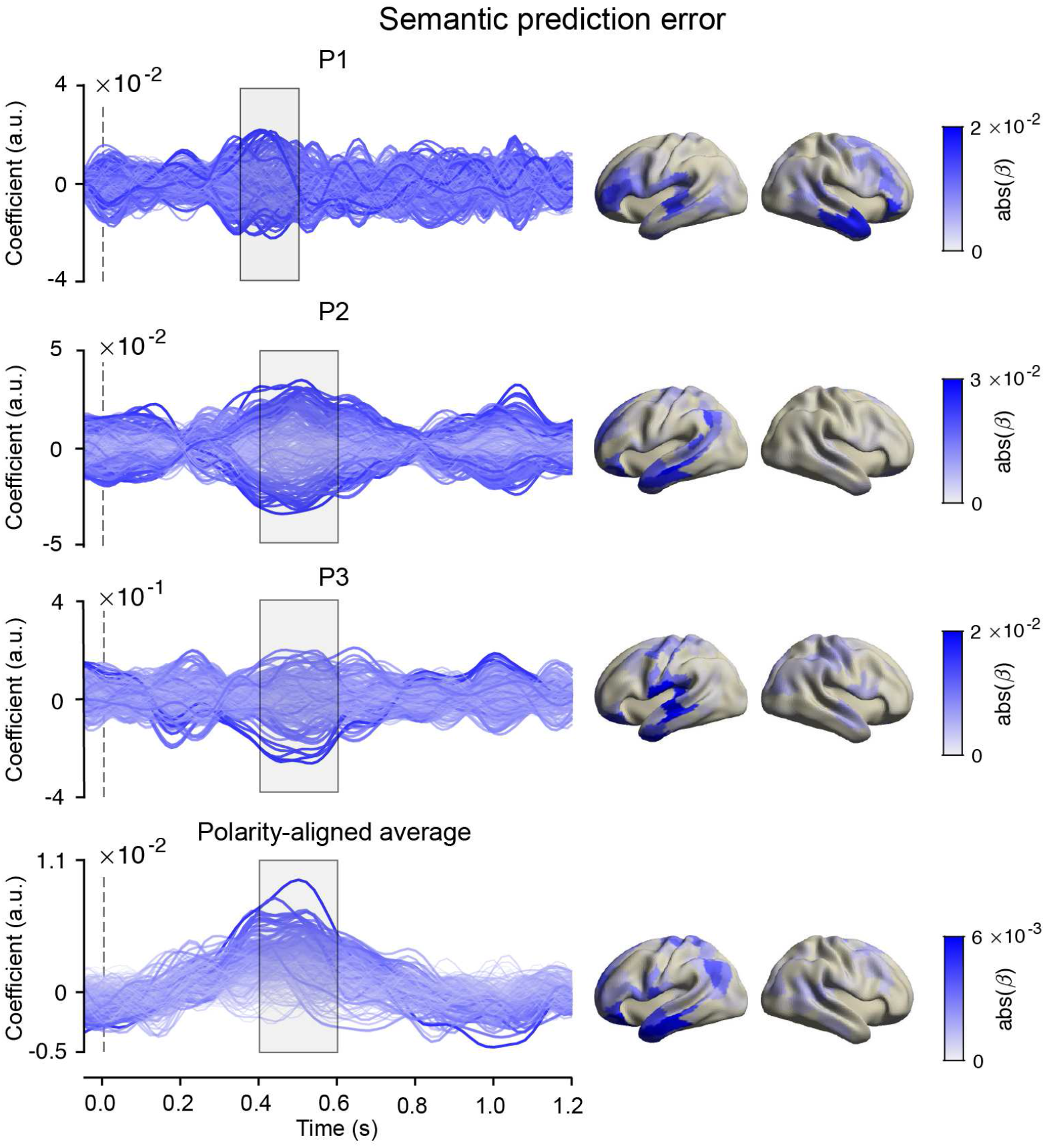
Coefficients for semantic prediction error from the integrated model (Figure S6) Left column: coefficients for each source for each individual in the MEG experiment, and the polarity-aligned average across participants. Right column: absolute value of the coefficients across the brain, averaged across the highlighted time-period.

**Figure S13.**
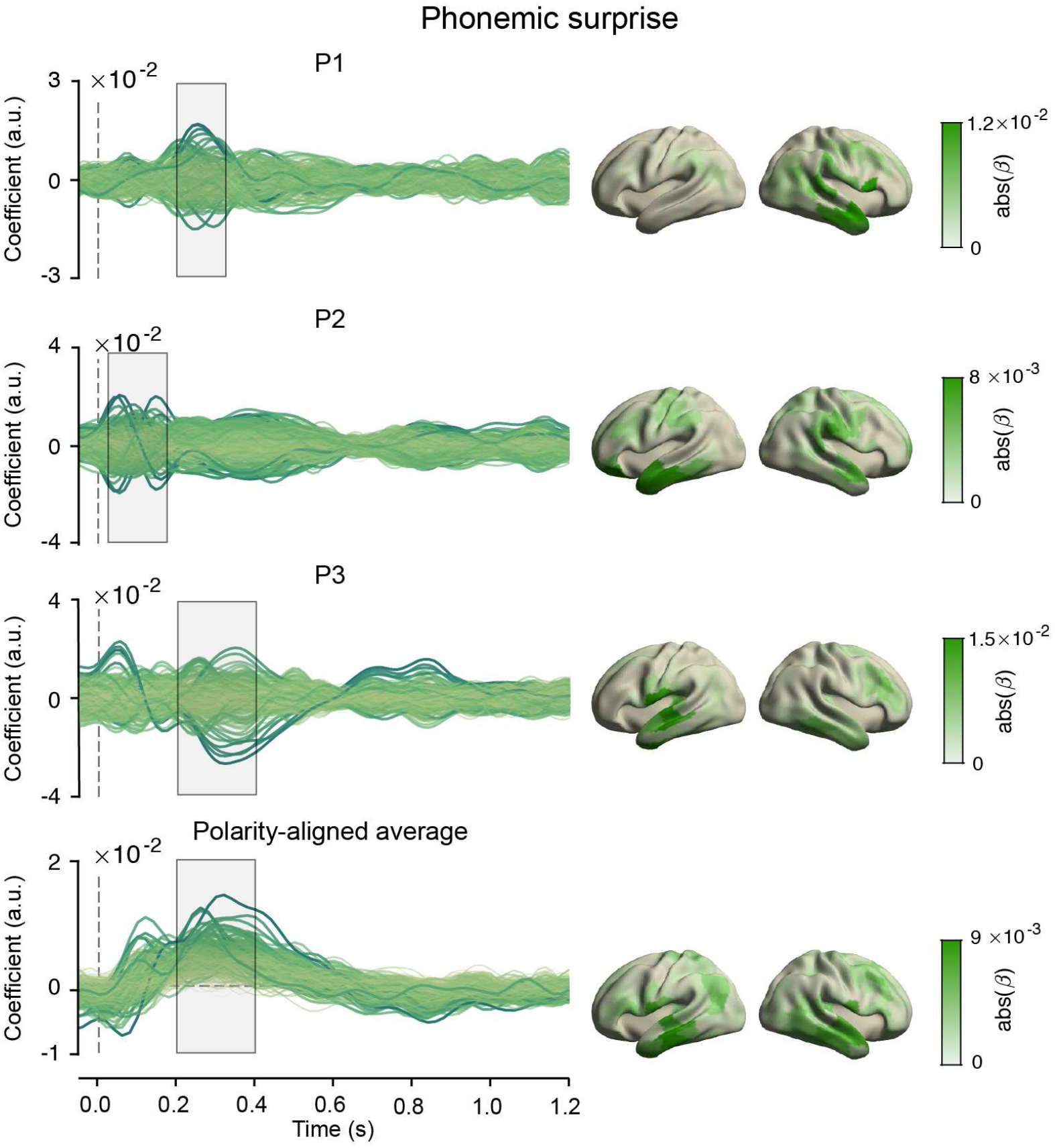
Coefficients for phonemic surprise from the integrated model (Figure S6) Left column: coefficients for each source for each individual in the MEG experiment, and the polarity-aligned average across participants. Right column: absolute value of the coefficients across the brain, averaged across the highlighted time-period..

**Figure S14.**
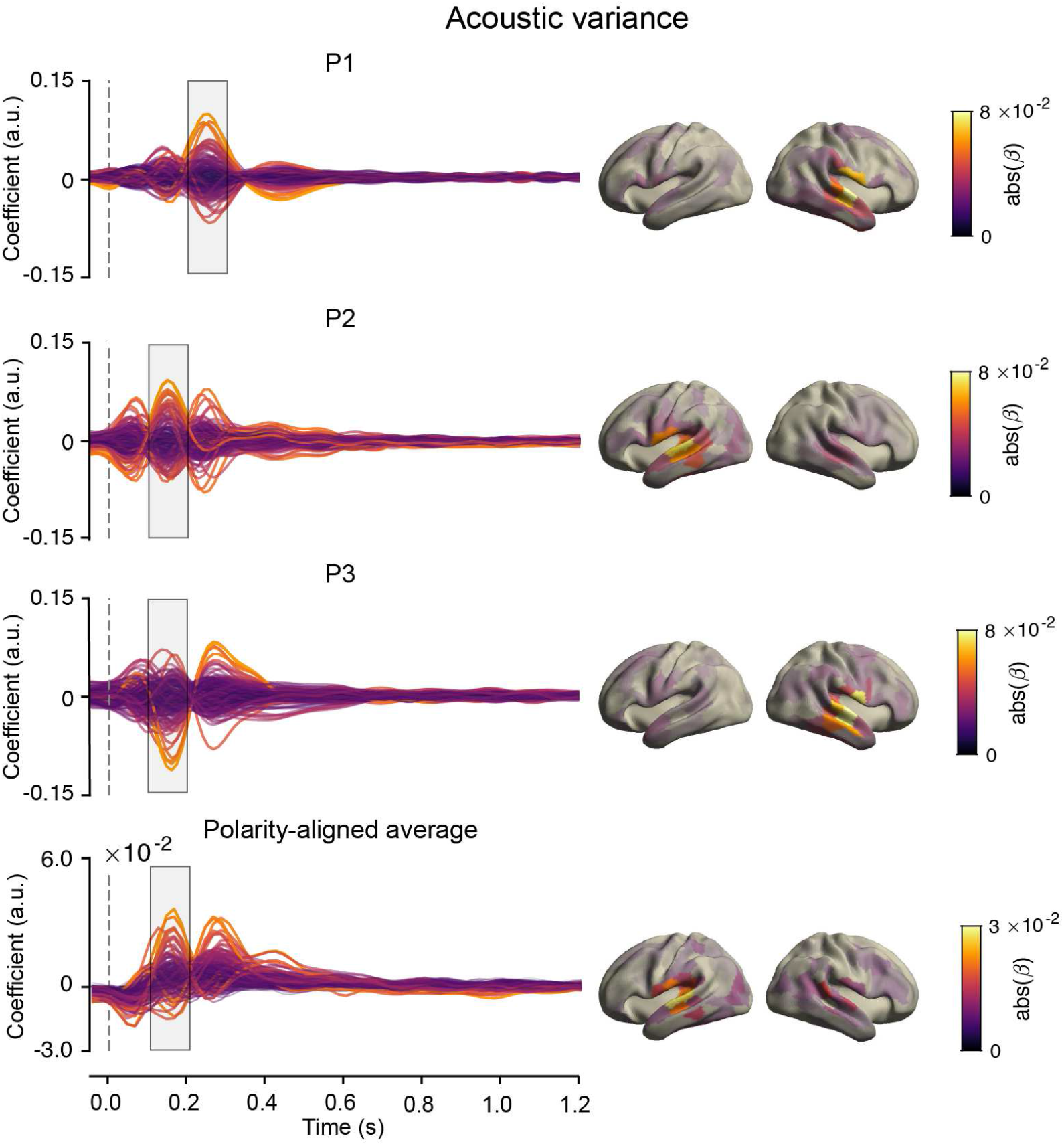
Coefficients for envelope variability from the integrated model (Figure S6) Left column: coefficients for each source for each individual in the MEG experiment, and the polarity-aligned average across participants. Right column: absolute value of the coefficients across the brain, averaged across the highlighted time-period.

**Figure S15.**
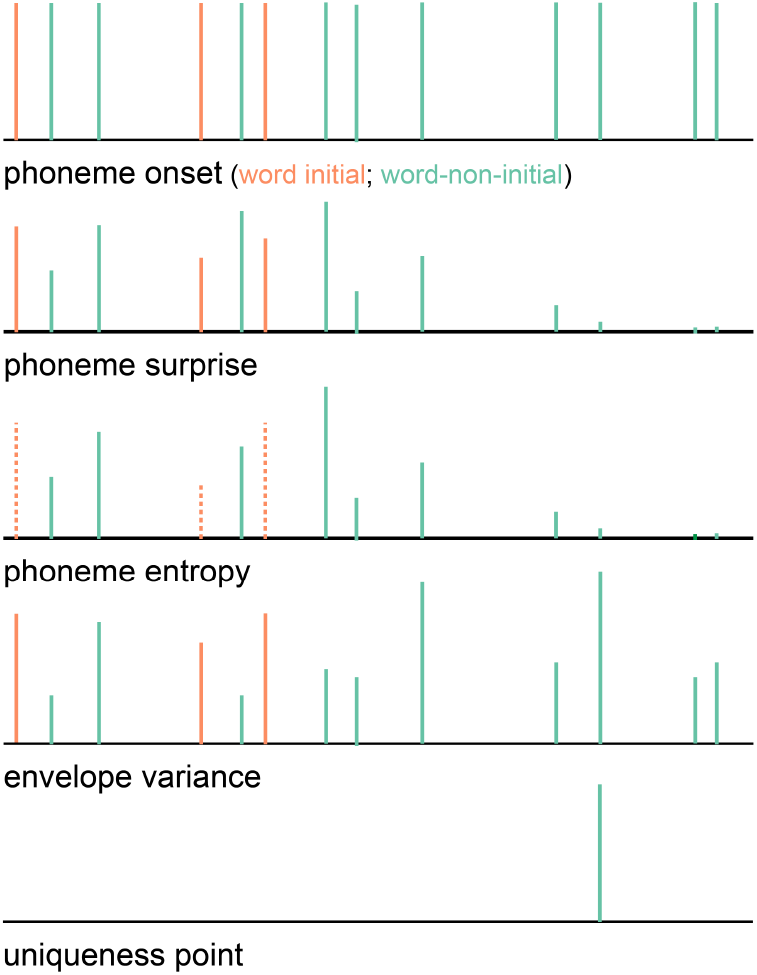
Regressors of the phoneme model. As indicated by the different colours, both the constants and covariates were modelled separately for word-initial and word-non-initial phonemes.

**Figure S16.**
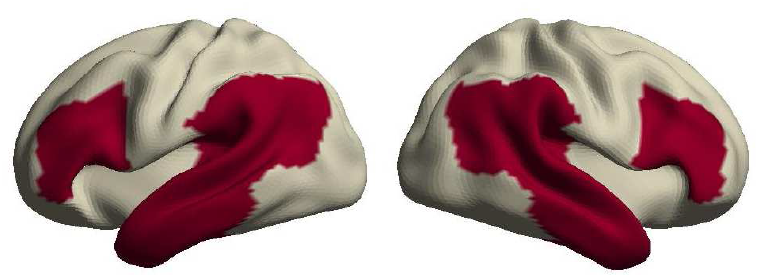
Language network definition. The language network was defined as temporal cortex plus temporo-parietal junction, and IFG and dorsolateral prefrontal cortex; all bilaterally. In terms of Brodmann areas this corresponded to 20, 21, 22, 38, 39, 40, 41, 42, 44, 45, 46 and 47, amounting to a total of 100 out of 370 cortical parcels.

